# RNA length and receptor usage define innate immune recognition across species

**DOI:** 10.64898/2026.05.21.726451

**Authors:** Julia Cieslicka, Karolina Drazkowska, Karolina Pianka, Wiktoria Szymanek, Rafal Tomecki, Charlotte Foret-Lucas, Dominik Cysewski, Romain Volmer, Pawel J. Sikorski

## Abstract

Recognition of viral double-stranded RNA (dsRNA) is a central trigger of innate antiviral immunity, yet the role of RNA length in shaping immune sensing across species remains poorly defined. Here, we identify RNA length as a key determinant of innate immune activation and demonstrate that this parameter is differentially tuned across vertebrate hosts. Using defined dsRNA molecules, we compared immune responses in human, duck, and chicken cells. Short 5’-triphosphorylated dsRNAs, including influenza-derived mini viral RNAs, robustly activated antiviral responses in human and duck cells but failed to do so in chicken cells. In contrast, longer dsRNAs induced immune activation across all species, revealing a species-specific threshold for dsRNA recognition. Mechanistically, this threshold is defined by differential engagement of RNA sensors. In human cells, sensing of short and intermediate dsRNA is strictly dependent on RIG-I, whereas in chicken cells, which lack RIG-I, MDA5 mediates dsRNA recognition and imposes a higher length requirement for activation. Consistent with this, duck cells, which retain RIG-I, respond to short dsRNA similarly to human cells. These differences in dsRNA sensing are accompanied by differential activation of downstream antiviral pathways, including PKR-dependent translational control and OAS/RNase L activation. Together, our findings identify RNA length as a fundamental parameter governing innate immune recognition and reveal that species-specific usage of RNA sensors constrains antiviral sensing, with implications for host susceptibility and virus-host interactions.

**Significance Statement:** Recognition of viral RNA is central to antiviral immunity, yet the rules governing which viral RNAs are sensed across species remain poorly understood. Here, we show that RNA length defines a species-specific threshold for innate immune recognition. Human and duck cells efficiently detect short viral double-stranded RNAs generated during influenza infection, whereas chicken cells respond only to substantially longer RNA duplexes. Mechanistically, this difference reflects the usage of distinct antiviral RNA sensors: RIG-I enables sensing of short viral RNAs, while MDA5 imposes a higher length requirement for activation. Our findings reveal that host species differ in the spectrum of viral RNA ligands accessible to immune detection, providing a framework for understanding species-specific antiviral responses and host susceptibility.

## Introduction

Innate immune recognition of viral RNA is a central component of host defense against RNA virus infections (1, 2). Among the molecular features sensed by host cells, double-stranded RNA (dsRNA) serves as a key trigger of antiviral responses. While the molecular pathways that detect dsRNA have been extensively characterized, the physical properties of RNA that determine immune recognition – and how these properties are interpreted across different host species – remain to be fully elucidated.

One such property is RNA length. Distinct antiviral sensors exhibit preferences for dsRNA molecules of different sizes: RIG-I preferentially recognizes short double-stranded RNAs bearing 5’-triphosphorylated ends (3, 4), whereas MDA5 (5-7) and downstream effectors such as PKR (8, 9) and the OAS/RNase L system (10-12) are typically activated by longer dsRNA species. This division of labor suggests that RNA length may function as a key parameter governing the engagement of antiviral pathways. However, whether RNA length imposes a general constraint on immune recognition across species, and how this relates to the usage of distinct RNA sensors in different hosts, remains unclear.

Recent studies have begun to reveal that immunostimulatory viral RNAs are not simply static products of canonical replication, but can arise from aberrant or noncanonical RNA synthesis driven by viral polymerase dynamics and RNA structure (13-16). In influenza A virus, during such processes short aberrant RNAs, including mini viral RNAs (mvRNAs), are generated as a consequence of dysregulated replication and transcription (13). These processes can be further modulated by host conditions such as temperature and the balance of viral proteins (17). mvRNAs are typically shorter than 100 nucleotides, form partially double-stranded structures with short duplex regions, and retain 5’-triphosphorylated ends (13) – features that promote recognition by RIG-I and underlie their potent immunostimulatory activity in mammalian cells. In addition, noncanonical transcription can generate complementary viral RNA species that further contribute to the formation of immunogenic RNA structures (18). Together, these observations indicate that the immunostimulatory RNA landscape is shaped not only by RNA length and sequence, but also by viral RNA biogenesis.

Host species differ in both the repertoire and functional tuning of dsRNA sensing pathways. In mammals and many avian species such as ducks, RIG-I serves as a primary sensor of influenza-derived RNA (19-21). In contrast, chickens lack RIG-I and rely on alternative receptors, including MDA5 and endosomal Toll-like receptors, to detect viral RNA (22-26). Notably, the absence of RIG-I is a conserved feature of *Galliformes* (landfowl), whereas it is retained in *Anseriformes* (waterfowl), including ducks (27). This difference has been proposed to contribute to variation in antiviral responses and susceptibility to influenza virus infection. In species lacking RIG-I, such as chickens, MDA5 is thought to contribute to viral RNA detection. MDA5 has been proposed to partially compensate for the absence of RIG-I in chickens. However, observations of increased susceptibility of chickens to several RNA viruses, including influenza and Newcastle disease virus (20, 28), suggest that this compensation may be incomplete. Whether the absence of RIG-I fully accounts for these species-specific differences remains unresolved. In particular, it is not known whether host species differ in the threshold at which dsRNA of varying lengths is recognized, whether such differences reflect the differential engagement of RNA sensors such as RIG-I and MDA5, and how this affects the detection of short viral RNAs generated during infection.

Here, we investigated how dsRNA length shapes innate immune activation across species. Using in vitro transcribed dsRNA molecules of defined lengths, we systematically compared immune responses in human, chicken, and duck cells. We find that short 5’-triphosphorylated dsRNAs robustly activate antiviral responses in human and duck cells but fail to do so in chicken cells, which require substantially longer dsRNA to trigger immune activation. This species-specific length threshold is defined by differential engagement of RNA sensors. In human cells, RIG-I mediates detection of short dsRNA, whereas in chicken cells – where MDA5 is the primary sensor – the absence of RIG-I imposes a higher effective length requirement. The ability of duck cells to respond to short dsRNA is consistent with their retention of RIG-I. Variation in dsRNA recognition across species is further reflected in downstream antiviral pathways, including PKR-dependent translational control and OAS/RNase L activation, and is associated with reduced responsiveness of chicken cells to influenza-derived short RNA species. Taken together, these results highlight the importance of RNA length as a pivotal factor in innate immune recognition and reveal that species-specific tuning of RNA sensing pathways imposes constraints on antiviral responses. This has implications for the host vulnerability to viral infections and virus-host interactions.

## Results

Influenza A virus generates a diverse set of RNA species during replication, including short aberrant transcripts that can engage cytoplasmic RNA sensors. Among these, mini viral RNAs (mvRNAs) are produced in substantial amounts during infection with highly pathogenic strains and act as potent agonists of RIG-I-mediated signaling in mammalian cells (13). Differences in the ability of host species to detect such RNA ligands may influence the efficiency of antiviral responses.

To address this, we examined how RNA length shapes innate immune activation across species by comparing responses in human cells and two avian systems – duck and chicken – which differ in their antiviral sensing capacity (20, 21). Using these models, we analyzed the immunogenicity of influenza-derived mvRNAs alongside synthetic dsRNA molecules of defined lengths.

### Species-specific differences in dsRNA sensing reveal a length-dependent threshold

To determine whether short viral RNAs are recognized differentially across species, we first examined the immunogenicity of an in vitro transcribed mini viral RNA (mvRNA)-like molecules in human, duck, and chicken cells. The mvRNA sequence was selected based on previously characterized influenza-derived RNAs shown to potently activate innate immune responses (13). The transcript used in this study corresponds to a 60-nucleotide RNA forming a duplex region of approximately 28 base pairs. Short 5’-triphosphorylated (5’-ppp) dsRNAs corresponding to mvRNA-like structures robustly induced antiviral responses in human and duck cells but failed to do so in chicken cells (Fig. 1).

**Figure 1.**
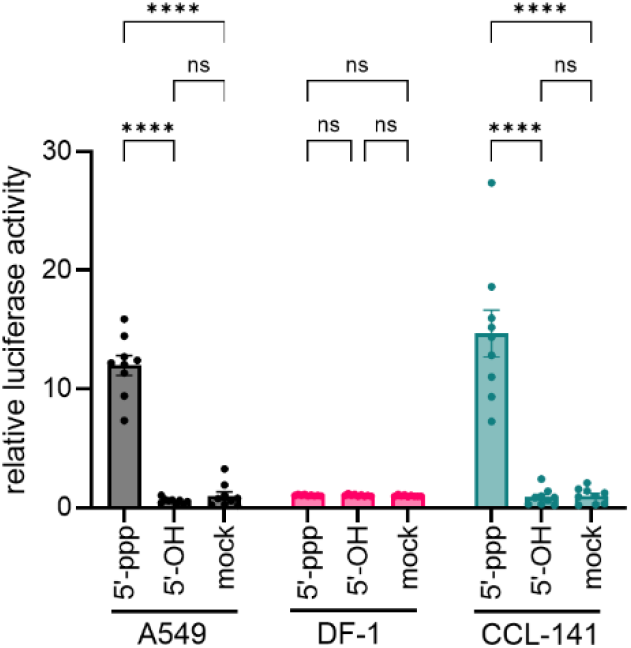
Species-specific recognition of mini viral RNA (mvRNA). Induction of antiviral response in human (A549), chicken (DF-1), and duck (CCL-141) cells following transfection with an in vitro transcribed mvRNA. The mvRNA used in this study is a 60 nt RNA forming an approximately 28 bp duplex and bearing a 5’-triphosphate (5’-ppp) or dephosphorylated (5’-OH) end. Luciferase activity driven by interferon-responsive promoters is shown. Cells were transfected with mvRNA for 24 h prior to measurement. Bars represent mean ± SEM from three independent biological replicates (each with three technical replicates), and individual points indicate technical replicates. Statistical significance: ****P < 0.0001; ns, not significant (one-way ANOVA with Tukey’s multiple comparisons test).

To confirm that immune activation depended on the presence of a 5’-triphosphate moiety, we compared the activity of 5’-ppp and dephosphorylated (5’-hydroxyl/OH) forms of the same RNA. Removal of the 5’-ppp group abolished immunogenicity in all tested cell types (Fig. 1), consistent with established requirements for recognition of short dsRNA (3, 4).

To determine whether this difference extends beyond mvRNA-like ligands and reflects a broader dependence on dsRNA length, we next examined immune responses to dsRNA molecules of increasing size. Using defined dsRNAs of 50, 200, 550, and 1600 bp (Fig. S1), we observed that both human and duck cells responded to short dsRNA in a manner strictly dependent on the presence of a 5’-triphosphate moiety, as dephosphorylated transcripts were non-immunogenic at these lengths (Fig. 2A).

**Figure 2.**
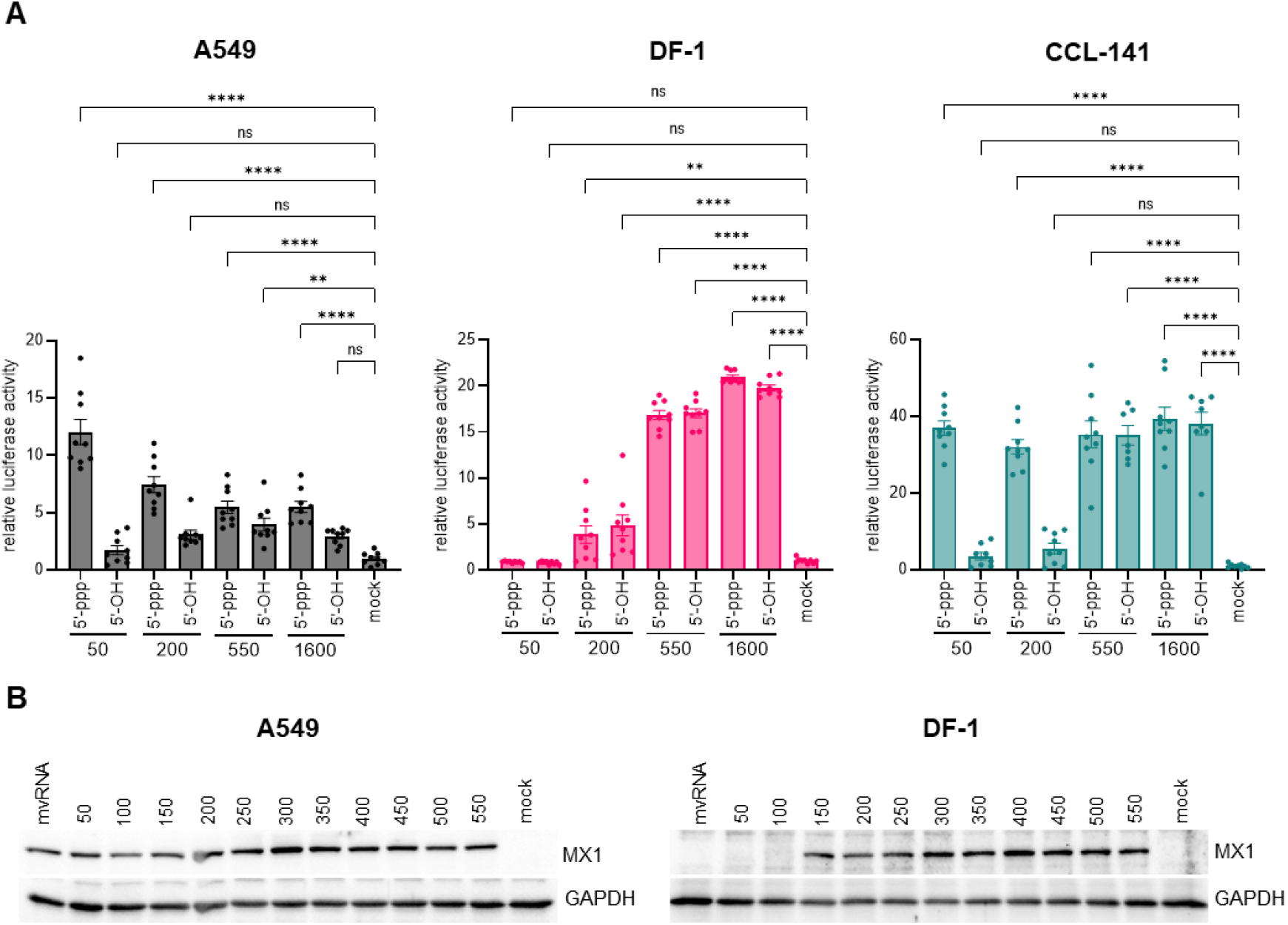
dsRNA length determines the threshold for innate immune activation across species. (A) Induction of antiviral response in human (A549), chicken (DF-1), and duck (CCL-141) cells following transfection with in vitro transcribed dsRNAs of defined lengths (50, 200, 550, and 1600 bp). dsRNAs carried either a 5’-triphosphate (5’-ppp) or a dephosphorylated (5’-OH) end. Luciferase activity driven by interferon-responsive promoters is shown. Cells were transfected with the indicated dsRNA for 24 h prior to measurement. Bars represent mean ± SEM from three independent biological replicates (each with three technical replicates), and individual points indicate technical replicates. Statistical significance is shown relative to mock-treated samples: **P < 0.01; ****P < 0.0001; ns, not significant (one-way ANOVA with Tukey’s multiple comparisons test). (B) Western blot analysis of MX1 protein expression in human and chicken cells following transfection with mvRNA and 5’-triphosphorylated dsRNAs of increasing lengths (50-550 bp).

At increased dsRNA lengths, however, species-specific differences became apparent. Duck cells exhibited robust responses to long dsRNA that were largely independent of 5’ end modification, with comparable activation observed for 5’-triphosphorylated and dephosphorylated duplexes. In contrast, human cells retained partial dependence on 5’ end chemistry, as dephosphorylated long dsRNA remained less immunogenic than their 5’-triphosphorylated counterparts.

Chicken cells displayed a distinct pattern, with short dsRNAs being largely non-immunogenic, an intermediate response emerging at approximately 200 bp, and robust activation observed with longer dsRNAs. Notably, responses to long dsRNA in chicken cells were largely independent of 5′ end modification, indicating a predominant reliance on RNA length rather than 5’ end features. Consistent with these observations, duck cells also displayed minimal dependence on 5’ end modification for long dsRNA sensing, whereas human cells retained partial dependence on 5’ end chemistry even for the longest dsRNA tested, highlighting a clear difference between avian and mammalian systems.

To more precisely define the length requirement for dsRNA sensing, we next analyzed immune activation across a more gradual range of dsRNA lengths between 50 and 550 bp. Based on initial luciferase reporter assays indicating a transition in immunogenicity at approximately 200 bp in chicken cells (Fig. 2A), we generated a series of in vitro transcribed dsRNAs starting at 50 bp and increasing in 50 bp increments up to 550 bp (Fig. S2). This length-dependent pattern of immune activation in chicken cells was further supported by western blot analysis of MX1 expression, an interferon-stimulated gene (ISG) product commonly used as a marker of antiviral activation. MX1 protein levels confirmed the absence of detectable protein induction at shorter dsRNA lengths and the onset of MX1 accumulation at approximately 150 bp in the DF-1 cell line. On the contrary, human cells showed robust MX1 expression across all tested lengths (Fig. 2B).

To assess whether these observations are applicable to physiologically relevant viral RNA species, we next examined the immunogenicity of short RNA fractions isolated from influenza-infected cells. Human A549 cells were infected with two distinct influenza A virus strains (highly pathogenic avian influenza virus H5N8 and the human-origin laboratory-adapted virus H1N1), both of which generate aberrant replication products, including mini viral RNAs (13). Total RNA was isolated from virus-infected cells as well as from dsRNA-transfected and mock-treated control cells. The RNA was then fractionated, and species shorter than 200 nucleotides were collected for further analysis (Fig. 3A, Fig. S3). Northern blot analysis confirmed that short cellular RNAs, including SNORD44 (63 nt), were preserved in the isolated fraction, indicating that fractionation did not disrupt the endogenous RNA population (Fig. 3B).

**Figure 3.**
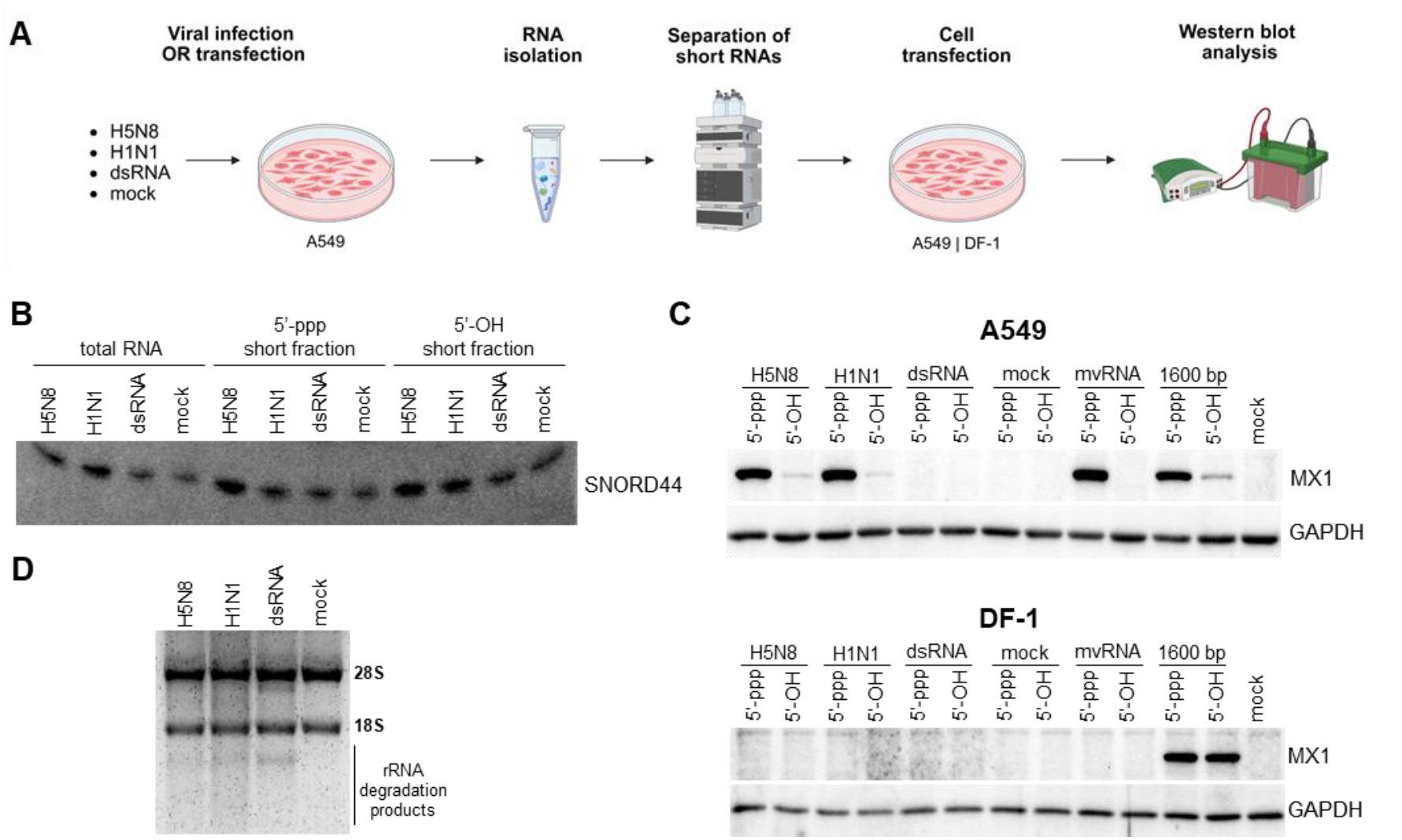
Short RNA fractions derived from influenza-infected cells are not immunogenic in chicken cells. (A) Schematic overview of RNA fractionation from virus (H5N8 or H1N1)-infected, dsRNA-treated, and mock-treated A549 cells using HPLC. Total RNA was fractionated, and RNA species shorter than 200 nt were collected. The short RNA fraction, either untreated or dephosphorylated with alkaline phosphatase, was used for transfection of human (A549) and chicken (DF-1) cells. (B) Northern blot analysis confirming the presence of endogenous short RNA, SNORD44 (63 nt), in the collected fraction. (C) Induction of antiviral response in human and chicken cells following transfection with the isolated short RNA fraction, assessed by western blot analysis of MX1 protein expression. (D) Analysis of total RNA isolated from virus-infected, dsRNA-treated, and mock-treated A549 cells. rRNA integrity was assessed in a denaturing agarose gel.

Transfection of the short RNA fraction from virus-infected cells induced robust antiviral responses in human cells, whereas no detectable activation was observed in chicken cells. Dephosphorylation of this fraction abolished its immunogenicity in human cells, indicating dependence on the 5’-triphosphorylated RNA species (Fig. 3C).

To determine whether immunogenicity could arise from nonspecific RNA degradation, we compared RNA derived from virus-infected cells with RNA isolated from cells treated with in vitro transcribed dsRNA. Both conditions resulted in comparable levels of ribosomal RNA (rRNA) degradation in human cells, indicative of similar RNase L activation (Fig. 3D). However, only the short RNA fraction from virus-infected cells elicited immune responses (Fig. 3C), indicating that immunogenicity is not a consequence of RNA degradation or the presence of nonspecific host-derived transcripts, but instead reflects the presence of specific viral RNA species.

Collectively, these results demonstrate that dsRNA sensing exhibits a species-dependent length threshold, with chicken cells requiring longer RNA duplexes for activation, whereas human and duck cells respond to a broader range of RNA lengths. Notably, despite retaining RIG-I, duck cells exhibited minimal dependence on 5’ end modification for sensing of long dsRNA, whereas human cells retained partial dependence on 5’ end chemistry.

### Species-specific receptor usage defines the length threshold for dsRNA sensing

To determine which RNA sensors underlie the observed species-specific differences in dsRNA recognition, we next examined the contribution of RIG-I and MDA5 to immune activation in human cells. A549 RIG-I knockout (KO) (29) and MDA5 KO cells (Fig. S4) were transfected with an in vitro transcribed mvRNA or with dsRNA molecules spanning a range of lengths from 50 to 550 bp. As shown in the preceding experiments, these RNA species robustly induced antiviral responses in wild-type cells. In contrast, no detectable immune activation was observed in RIG-I KO cells across all tested RNA ligands, whereas MDA5 KO cells displayed responses comparable to those previously observed in wild-type cells (Fig. 4). These results demonstrate that sensing of short and intermediate dsRNA, including mvRNA, in human cells is strictly dependent on RIG-I and does not require MDA5.

**Figure 4.**
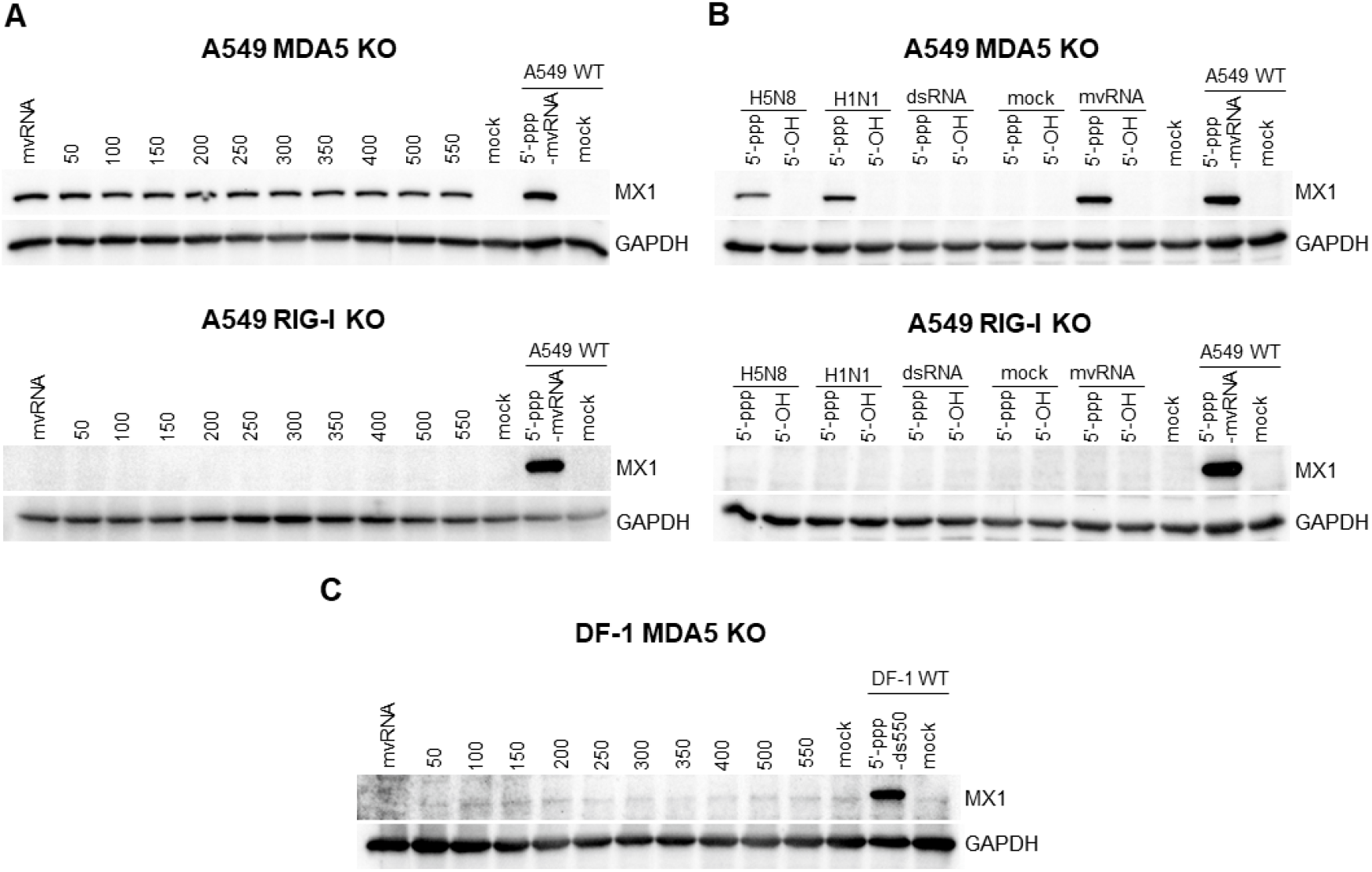
Species-specific receptor usage defines the length threshold for dsRNA sensing. (A) Western blot analysis of antiviral responses assessed by MX1 expression in human (A549) cells lacking RIG-I or MDA5 following transfection with 5’-triphosphorylated in vitro transcribed mvRNA and 5’-triphosphorylated dsRNAs of increasing lengths. Lanes labeled “A549 WT” represent wild-type A549 cells transfected with 5’-ppp mvRNA and are included as positive controls for immune activation. (B) Induction of antiviral response in human cells lacking RIG-I or MDA5 following transfection with the isolated short RNA fraction, assessed by western blot analysis of MX1 expression. Lanes labeled “A549 WT” represent wild-type A549 cells transfected with with 5’-ppp mvRNA and are included as positive controls for immune activation. (C) Induction of antiviral response in chicken cells lacking MDA5 following transfection with 5’-triphosphorylated in vitro transcribed mvRNA and 5’-triphosphorylated dsRNAs of increasing lengths, assessed by western blot analysis of MX1 expression. Lanes labeled “DF-1 WT” represent wild-type DF-1 cells transfected with 5’-ppp 550 bp dsRNA and are included as positive controls for immune activation.

To assess whether this requirement extends to physiologically relevant viral RNA species, we next analyzed the immunogenicity of short RNA fractions isolated from influenza-infected cells in the same genetic backgrounds. Consistent with the results obtained using synthetic RNA ligands, transfection with the short RNA fraction failed to induce detectable responses in RIG-I KO cells, whereas MDA5 KO cells exhibited robust activation comparable to that observed in wild-type cells (Fig. 4). These findings indicate that RIG-I is the primary sensor responsible for detection of short viral RNA species generated during influenza infection.

We next examined the contribution of MDA5 to dsRNA sensing in chicken cells. DF-1 MDA5 KO cells (Fig. S5) were transfected with in vitro transcribed mvRNA as well as dsRNA molecules spanning 50 to 550 bp. In contrast to the robust, length-dependent induction of MX1 observed in wild-type DF-1 cells in the preceding experiments, MDA5 deficiency resulted in a complete loss of MX1 protein expression across all tested RNA ligands (Fig. 4), demonstrating that MDA5 is required for dsRNA-mediated immune activation in chicken cells.

Together, these results reveal that the species-specific usage of RNA sensors defines the effective length threshold for dsRNA recognition. In human cells, RIG-I enables detection of short and intermediate dsRNA species, including viral RNA fragments such as mvRNAs. In contrast, chicken cells rely on MDA5, which preferentially responds to longer dsRNA, thereby imposing a higher length requirement for activation and limiting responsiveness to shorter RNA ligands.

### Length-dependent dsRNA sensing differentially activates PKR and translational control across species

Having established that species-specific receptor usage defines the length threshold for dsRNA sensing, we next asked how these differences influence activation of downstream antiviral pathways, focusing on PKR-mediated translational control (30).

For the sake of clarity, in the following experiments, “short dsRNA” refers to mvRNA-like duplex and “long dsRNA” to the 1600 bp duplex, and this terminology is used consistently throughout the remainder of the manuscript.

As a preliminary step, we sought to ascertain the reliability of the eIF2α phosphorylation and puromycin incorporation assays in reflecting translational inhibition in the tested cell lines. Treatment with sodium arsenite, a known inducer of translational arrest, resulted in robust phosphorylation of eIF2α and a marked reduction in puromycin incorporation in all cell types, confirming that this experimental setup is suitable for monitoring translational responses in both human and avian cells (Fig. S6).

We next examined PKR activation following transfection with dsRNA of different lengths and 5’ end configurations. Short dsRNA failed to induce detectable eIF2α phosphorylation in any of the tested cell lines, regardless of 5’ modification. In contrast, long dsRNA triggered strong eIF2α phosphorylation in human and chicken cells irrespective of its 5’ end modification, whereas duck cells exhibited a consistently weaker response (Fig. 5A).

**Figure 5.**
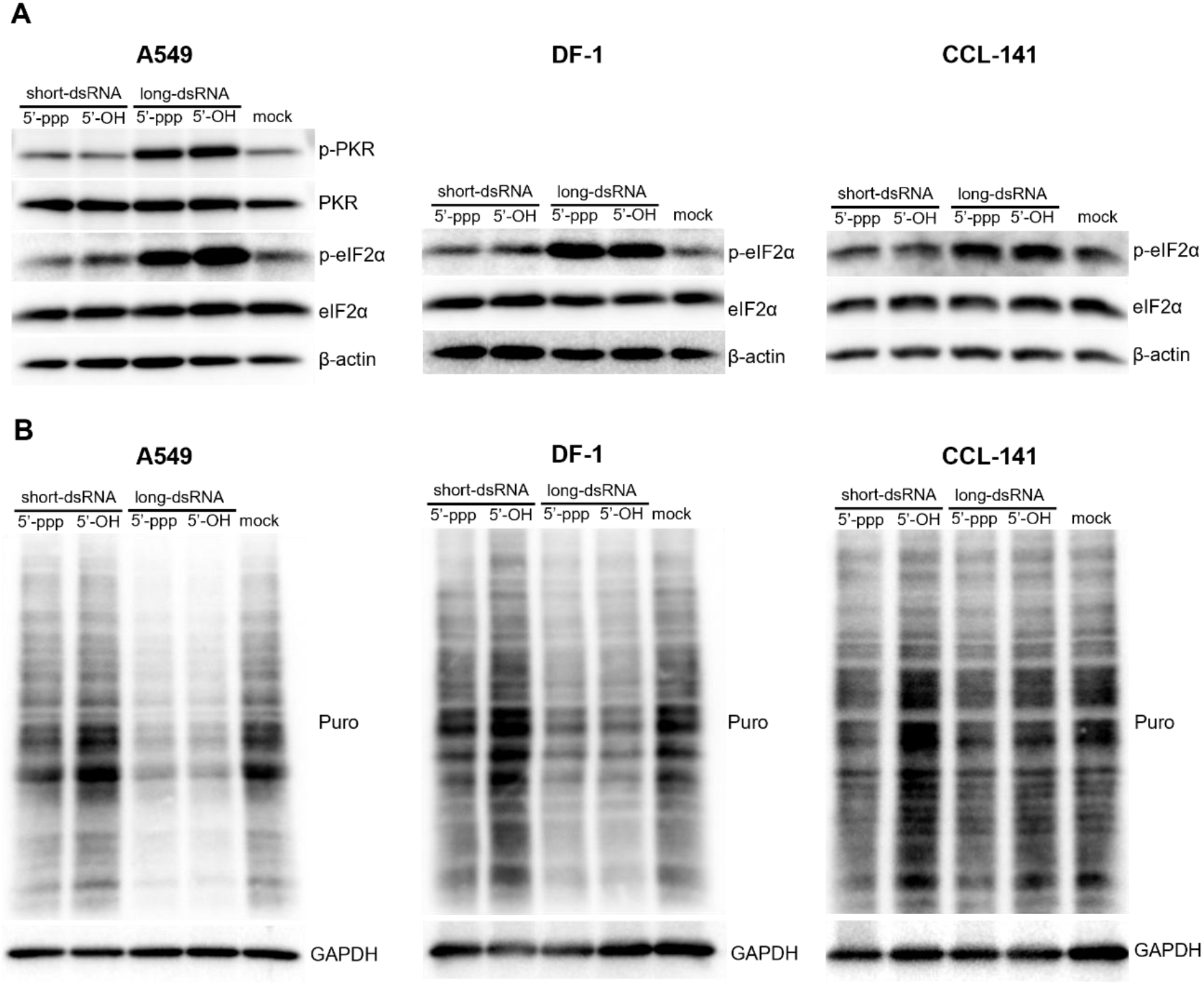
Length-dependent activation of PKR and translational control across species. (A) Western blot analysis of PKR and eIF2α phosphorylation in human (A549), chicken (DF-1), and duck (CCL-141) cells following transfection with short (mvRNA) or long (1600 bp) dsRNA carrying either a 5’-triphosphate (5’-ppp) or a dephosphorylated (5’-OH) end. Human and chicken cells were transfected for 5 h, whereas duck cells were transfected for 7.5 h prior to harvesting. PKR phosphorylation was assessed in human samples. (B) Puromycin incorporation assay to assess global translation in human, chicken, and duck cells following transfection with short or long dsRNA. Reduced puromycin signal indicates translational inhibition.

Consistent with these findings, puromycin incorporation assays revealed that long dsRNA induced pronounced translational inhibition in human and chicken cells, while duck cells displayed only partial reduction in protein synthesis under the same conditions. Short dsRNA demonstrated a negligible impact on translation across the examined systems (Fig. 5B).

Together, these results indicate that efficient activation of PKR and induction of translational shutdown require long dsRNA. Notably, despite robust sensing of short dsRNA in human cells mediated by RIG-I, these ligands are insufficient to trigger PKR activation, highlighting a functional separation between RNA sensing and translational control pathways. Furthermore, the magnitude of PKR activation varies among studied species, with reduced responses observed in duck cells compared to human and chicken cells.

### Long dsRNA preferentially recruits antiviral RNA-binding proteins across species

Having established that dsRNA length determines both immunogenicity and PKR activation, and that sensing of short and intermediate dsRNA is mediated by distinct RNA receptors across species, we next sought to determine whether these differences are reflected in the recruitment of cellular RNA-binding proteins.

Biotinylated short and long dsRNAs, carrying either a 5’-ppp group or a dephosphorylated (5’-OH) end (Fig. S7), were used in pull-down assays with lysates from human, chicken, and duck cells. To account for potential effects of antiviral priming, experiments were performed using lysates from both stimulated and mock-treated cells, and comparable binding patterns were observed under both conditions. Bound proteins were subsequently identified by mass spectrometry. It is important to note that this approach primarily captures stable RNA-protein interactions formed under these conditions and may not detect more transient or context-dependent interactions.

Across all species, long dsRNA molecules were associated with a broad set of RNA-binding proteins, whereas short dsRNA consistently showed minimal or no specific protein enrichment, irrespective of 5’ end modification (Fig. 6). Notably, this occurred despite the robust immunogenicity of 5’-ppp short dsRNA in human and duck cells, indicating that efficient immune activation in these systems can be achieved in the absence of stable or abundant protein binding detectable by this approach.

**Figure 6.**
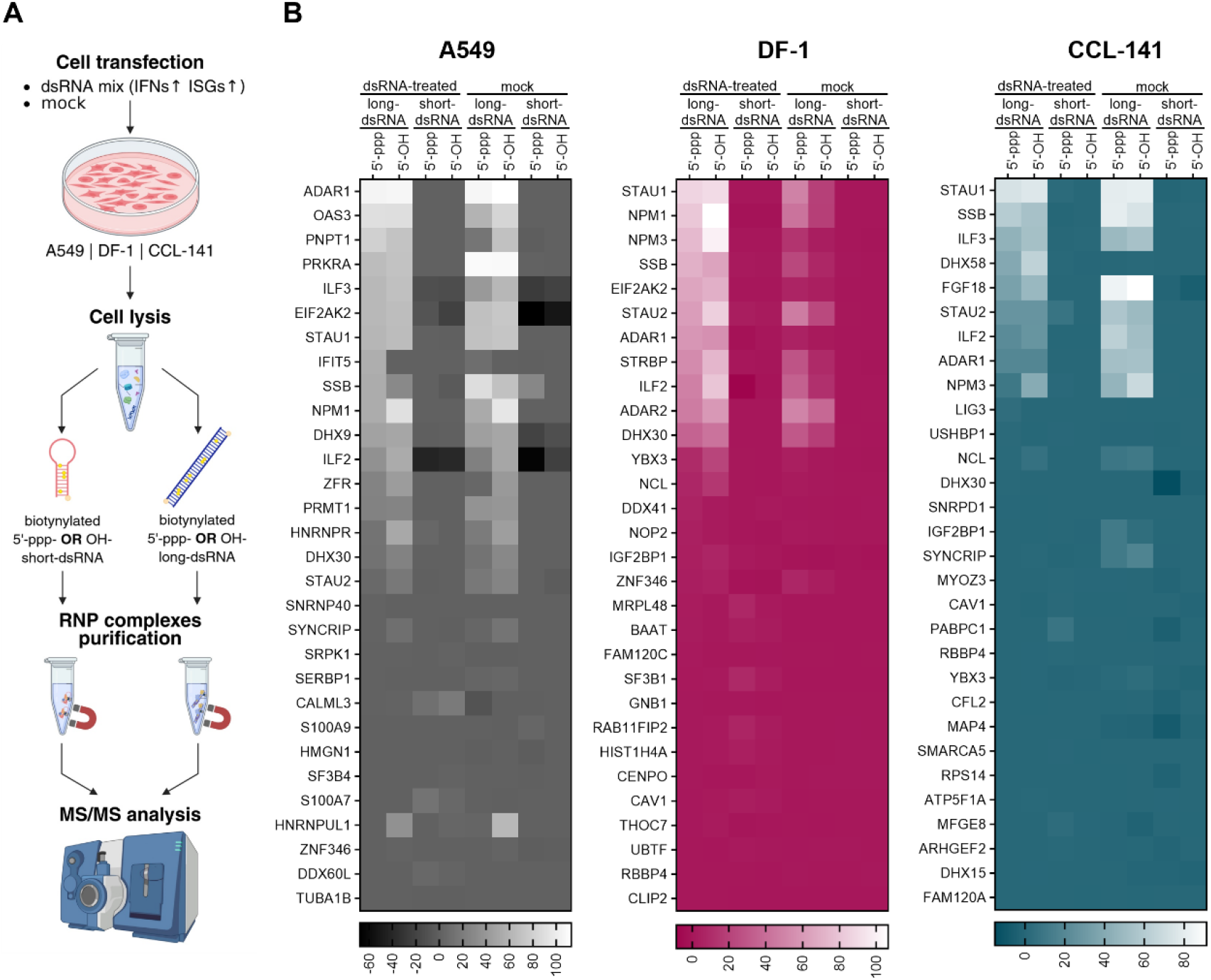
Length-dependent recruitment of RNA-binding proteins across species. (A) Schematic overview of the pull-down assay. Human (A549), chicken (DF-1), and duck (CCL-141) cells were transfected with a mixture of 5’-triphosphorylated short (mvRNA) and long (1600 bp) dsRNA or mock-treated prior to lysis. Cell lysates were incubated with biotinylated short or long dsRNA carrying either a 5’-triphosphate (5’-ppp) or a dephosphorylated (5’-OH) end. RNA-protein complexes were isolated using streptavidin beads and analyzed by mass spectrometry (MS/MS). (B) Heatmaps showing enrichment of RNA-binding proteins in pull-down assays from human, chicken, and duck cell lysates. Protein enrichment is displayed as a composite score calculated as log2(FC) multiplied by -log10(p-value), relative to beads-only control samples. Color scales indicate relative protein enrichment. The top 30 enriched proteins are shown; full datasets are provided in the supporting information (Fig. S11).

Among the proteins enriched on long dsRNA, PKR (EIF2AK2) was strongly detected in pull-downs from human and chicken cell lysates, whereas its enrichment was less pronounced in duck cells (Fig. 6 and Fig. S8-S11), consistent with the differences in PKR activation and translational inhibition observed across species (Fig. 5).

In contrast, short dsRNA failed to efficiently recruit PKR or other detectable dsRNA-binding proteins in all tested systems. Together with the receptor dependency identified above, these findings suggest that recognition of short dsRNA, particularly in human cells, likely involves transient or low-abundance interactions – such as those mediated by RIG-I – that are not efficiently captured under these experimental conditions.

Together, these results indicate that dsRNA length critically influences the recruitment of cellular RNA-binding proteins. Preferential engagement of antiviral effectors by longer dsRNA, rather than differences in 5’ end chemistry or cellular activation state, likely contributes to the observed species-specific differences in downstream antiviral responses.

### Differential engagement of the OAS/RNase L pathway across species

To determine whether length-dependent differences in dsRNA sensing extend to additional antiviral pathways, we next examined activation of the OAS/RNase L pathway across species (31).

Activation of this pathway was assessed by monitoring rRNA degradation following transfection of cells with short and long dsRNA carrying either a 5’-ppp or a dephosphorylated (5’-OH) end. In human cells, transfection with long dsRNA resulted in clear rRNA degradation, indicative of robust RNase L activation, whereas short dsRNA did not induce detectable rRNA cleavage (Fig. 7). In contrast, no evidence of rRNA degradation was observed in chicken or duck cells under the same conditions. This was not due to inefficient stimulation or transfection, as the same dsRNA conditions induced robust immune activation in avian cells (Fig. 2).

**Figure 7.**
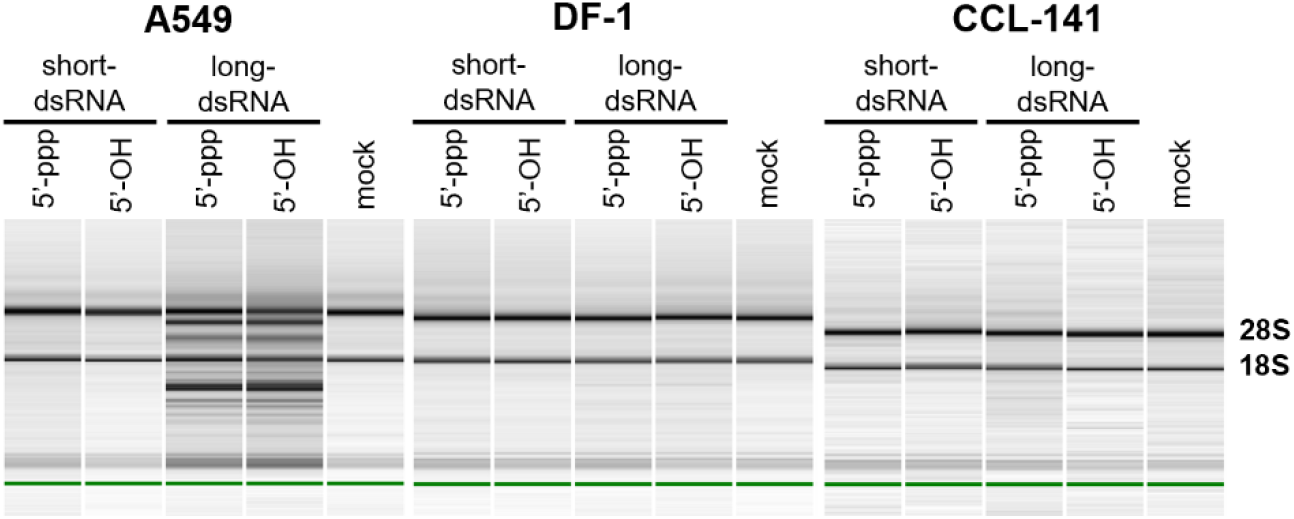
Activation of the OAS/RNase L pathway is observed in human but not avian cells. Analysis of total RNA integrity in human (A549), chicken (DF-1), and duck (CCL-141) cells following transfection with short (mvRNA) or long (1600 bp) dsRNA carrying either a 5’-triphosphate (5’-ppp) or a dephosphorylated (5’-OH) end. RNA profiles were assessed using Agilent 2100 Bioanalyzer.

These results indicate that the activation of the OAS/RNase L pathway is restricted to human cells and requires long dsRNA, further reinforcing the length-dependent nature of antiviral pathway engagement. Notably, this pattern mirrors the preferential activation of PKR by long dsRNA and highlights a functional separation between RNA sensing and downstream effector pathways.

Consistent with these observations, OAS3 was selectively enriched in pull-down assays from human cell lysates, whereas no enrichment of OASL was detected in avian samples (Fig. 6). Notably, avian species encode OASL rather than the multi-member OAS1-3 family present in humans (32-34), which may influence the efficiency of dsRNA-dependent activation of the OAS/RNase L pathway. Overall, these findings indicate that, in contrast to human cells, avian systems do not efficiently engage the OAS/RNase L pathway in response to dsRNA, revealing an additional layer of species-specific divergence in antiviral responses.

## Discussion

Recognition of viral RNA by innate immune sensors is a fundamental determinant of antiviral defense (1, 2), yet the principles governing how physical properties of RNA shape this process across species have remained incompletely understood. Here, we demonstrate that dsRNA length defines a critical threshold for innate immune activation that is differentially tuned across vertebrate hosts. Importantly, we show that this threshold is not solely an intrinsic property of RNA itself but is determined by the repertoire of RNA sensors expressed in a given species.

Recent studies provide important context for interpreting these findings by linking the generation of immunostimulatory viral RNAs to the underlying mechanisms of viral RNA synthesis. Influenza virus replication is inherently error-prone and can give rise to a spectrum of aberrant RNA products, including mini viral RNAs (mvRNAs), through polymerase stalling and template-dependent RNA structures (13-15, 18). The production of these RNAs is further influenced by host conditions, such as body temperature and the balance of viral proteins, which modulate polymerase processivity and favor the accumulation of short, noncanonical RNA species (17). In addition, emerging evidence indicates that aberrant transcription can generate complementary RNA products, potentially expanding the repertoire of immunostimulatory RNA structures present in infected cells (18).

Within this framework, RNA length can be viewed as a determinant that reflects how host cells filter a heterogeneous pool of viral RNA products generated during infection. While influenza-derived short RNAs such as mvRNAs fall within the recognition range of RIG-I, they remain below the effective activation threshold of MDA5. As a result, species that lack RIG-I, such as chickens, are likely unable to efficiently detect early, short RNA products of viral replication, whereas species that retain RIG-I can respond to these ligands. This provides a mechanistic basis for how differences in receptor usage translate into species-specific sensitivity to distinct classes of viral RNA.

Our results identify differential usage of RIG-I and MDA5 as the key mechanism underlying species-specific differences in dsRNA sensing. In human cells, RIG-I enables robust detection of short and intermediate dsRNA species, including influenza-derived mvRNAs. This observation is consistent with previous studies indicating that RIG-I plays a more prominent role than MDA5 in sensing short and intermediate dsRNA in human cells (3-7). Our findings extend this concept by demonstrating that RIG-I is not only sufficient but strictly required for detection of dsRNA ligands within this size range. In contrast, chicken cells, which lack RIG-I (27), rely on MDA5, resulting in a higher length requirement for immune activation. This finding provides a mechanistic explanation for the limited responsiveness of chicken cells to short dsRNA ligands and establishes receptor usage as a factor shaping the effective range of RNA lengths that can be sensed.

These observations also help clarify longstanding questions regarding the relationship between RIG-I and MDA5. While MDA5 has been proposed to compensate for the absence of RIG-I in species such as chickens (23, 25, 27), our data indicate that this compensation is incomplete, at least with respect to the detection of short RNA ligands. Rather than fully substituting for RIG-I, MDA5 appears to impose distinct constraints on RNA recognition that favor longer duplex substrates. In this context, previous studies have reported that chicken MDA5 can respond to relatively short dsRNA compared to its mammalian counterpart, primarily based on experiments using poly(I:C) species in the range of ∼0.2-1 kb (25). However, these RNAs remain relatively long and fully duplexed, and therefore do not address whether MDA5 efficiently recognizes short duplexes in the tens-of-base-pairs range. Structural studies further support a model in which MDA5 engages duplex RNA through cooperative filament formation along the RNA backbone, a mechanism that inherently favors extended dsRNA substrates (35-38). In contrast, our data reveal a clear functional threshold for dsRNA sensing in chicken cells, with efficient immune activation requiring duplexes of at least ∼150–200 bp under the tested conditions. This supports a model in which, in a cellular context, MDA5 imposes more stringent requirements on ligand length than previously appreciated. This limited responsiveness to shorter RNA ligands may contribute, at least in part, to the increased susceptibility of chickens to RNA virus infections compared to species such as ducks (20, 28), which retain RIG-I and are capable of efficiently detecting short 5’-triphosphorylated RNA species.

Taken together, these findings have important implications for host-virus interactions. Influenza A virus produces short, 5’-triphosphorylated RNA species that are potent activators of RIG-I in mammalian cells. These mini viral RNAs are typically shorter than 100 nucleotides and form duplex regions of less than ∼50 base pairs, placing them below the effective length threshold required for MDA5 activation. Our findings suggest that the inability of chicken cells to detect such RNAs may limit early antiviral signaling, potentially contributing to differences in susceptibility to infection observed between avian species. In this context, the presence or absence of RIG-I (27) – retained in *Anseriformes* but lost in *Galliformes* – may represent an important evolutionary determinant of antiviral defense strategies.

Beyond initial sensing, we find that dsRNA length also governs activation of downstream antiviral pathways. While short dsRNA efficiently triggers RIG-I-dependent signaling in human cells, it is insufficient to robustly activate PKR or induce translational shutdown. This is consistent with the known mechanism of PKR activation, which requires binding to dsRNA followed by RNA-mediated dimerization and autophosphorylation (8, 9, 39), imposing a strict dependence on RNA length. Short duplexes are therefore below the threshold required to support productive PKR activation, whereas longer dsRNA species provide an adequate platform for PKR dimerization and signaling. Accordingly, long dsRNA robustly engages PKR across species, leading to inhibition of protein synthesis. PKR has been reported to be upregulated in response to viral infection in both duck and chicken systems, including during highly pathogenic avian influenza virus infection (40-44), indicating that this pathway is conserved and responsive in avian hosts. However, our data suggest that the magnitude and cellular consequences of PKR activation are modulated by dsRNA length and may differ between species. In particular, the comparatively weaker PKR activation observed in duck cells, despite their ability to sense short dsRNA, highlights a functional uncoupling between RNA sensing and downstream effector activation.

Similarly, activation of the OAS/RNase L pathway was observed only in human cells and required long dsRNA, consistent with selective enrichment of OAS3 in pull-down assays. In mammals, efficient activation of this pathway depends on high-affinity dsRNA sensors such as OAS3 (12, 29). In contrast, avian species typically encode OASL (32-34), which contains a single OAS-like domain and may differ in its efficiency of dsRNA-dependent activation. Although avian OASL retains catalytic residues required for 2’-5’ oligoadenylate synthesis (32-34), it may not generate sufficient levels of 2’-5’ oligoadenylates to reach the activation threshold of RNase L under the conditions tested here. Accordingly, we did not observe detectable RNase L activation in avian cells following dsRNA stimulation. Importantly, the lack of detectable RNase L activation does not exclude antiviral functions of OASL, which have been reported to operate independently of canonical 2’-5’ oligoadenylate synthesis and RNase L activation (33, 45-47). Thus, OASL-mediated antiviral activity in avian cells may occur through alternative mechanisms.

Our study also provides insight into the molecular basis of dsRNA recognition. Despite their immunogenicity, short dsRNA ligands showed minimal recruitment of RNA-binding proteins in pull-down assays, suggesting that their detection may rely on transient or low-abundance interactions that are not readily captured by biochemical approaches. In contrast, long dsRNA formed stable complexes with multiple antiviral effectors, including PKR and OAS3. These observations highlight the importance of considering both binding stability and functional outcomes when interpreting RNA-protein interactions.

Together, our findings establish RNA length as a central determinant of innate immune recognition and demonstrate that species-specific usage of RNA sensors defines the effective range of viral RNA ligands that can be detected. Importantly, our results place this principle in the broader context of viral RNA biogenesis, suggesting that the pool of immunostimulatory RNA species is shaped by the dynamics of viral replication and transcription. Within this framework, RNA length emerges not merely as an intrinsic property of viral ligands, but also as a parameter that determines which products of viral RNA synthesis become accessible to host defense pathways. Differences in sensor repertoires across species therefore translate into distinct windows of sensitivity to viral RNA, with important consequences for the timing and magnitude of antiviral responses. These findings provide a unifying framework for understanding how physical properties of RNA and species-specific receptor usage jointly shape innate immune recognition, with implications for host susceptibility, viral pathogenesis, and the evolution of antiviral defense mechanisms.

## Materials and Methods

### Cell lines

Human lung epithelial A549 cells (CCL-185) and chicken embryo fibroblast DF-1 cells (CRL-3586) were obtained from ATCC. A549 RIG-I knockout (KO) cells were generated previously (29), whereas A549 MDA5 KO and DF-1 MDA5 KO cell lines were generated in this study. A549 and DF-1 cells were maintained at 37°C in a humidified atmosphere with 5% CO_2_ in Dulbecco’s Modified Eagle Medium (DMEM; Gibco 21969035) supplemented with 10% (v/v) heat-inactivated fetal bovine serum (FBS; Sigma-Aldrich F9665), 1% (v/v) GlutaMAX (Gibco 35050061), and 1% (v/v) penicillin-streptomycin (Gibco P4333). Duck embryo fibroblast cells (CCL-141; ATCC) were maintained under the same conditions in Minimum Essential Medium (MEM; Gibco 11095080) supplemented with 10% (v/v) heat-inactivated FBS, 1% (v/v) penicillin/streptomycin, 2 mM L-glutamine (Gibco 25030081), 1 mM sodium pyruvate (Gibco 11360070), 1.5 g/L sodium bicarbonate (Gibco 25080094), and 1x non-essential amino acids (Gibco 11140035).

### Viruses

The highly pathogenic avian influenza A virus H5N8 was generated by reverse genetics as described previously, using the eight genomic segments of the A/mallard duck/France/171201g/2017 (H5N8) field isolate (48). These viruses were approved by the French biotechnology ethics committee (Comité d’expertise des utilisations confinées d’OGM) and were handled exclusively in biosafety level 3 laboratories. The human-origin influenza A virus H1N1 was also generated by reverse genetics using a set of eight pHW2000 plasmids (49) containing the genomic segments of A/WSN/1933 (H1N1) (50).

### Antibodies

The following primary antibodies were used for western blotting at a dilution of 1:1000: mouse anti-β-actin (Santa Cruz Biotechnology sc-47778, RRID: AB_626632), rabbit anti-eIF2α (Cell Signaling 9722S, RRID:AB_2230924), rabbit anti-phospho-eIF2α (Ser51) (Cell Signaling 9721S, RRID:AB_330951), rabbit anti-GAPDH (Novus NB300-327, RRID:AB_10001915), rabbit anti-MDA5 (ABclonal A2419, RRID: AB_3674625), mouse anti-MX1 (Merck MABF938, RRID:AB_2885181), rabbit anti-PKR (Cell Signaling 12297S, RRID:AB_2665515), rabbit anti-phospho-PKR (Thr444) (ABclonal AP1134, RRID: AB_2864002), mouse anti-Puromycin (Sigma Aldrich MABE343, RRID:AB_2566826). Horseradish peroxidase (HRP)-conjugated secondary antibodies were used at a dilution of 1:10000: goat anti-Mouse IgG (H+L) Secondary Antibody HRP (Invitrogen 62-6520, RRID:AB_2533947) and goat anti-Rabbit IgG (H+L) Secondary Antibody HRP (Invitrogen 65-6120, RRID:AB_2533967).

### Plasmids

To generate a plasmid encoding mini viral RNA (mvRNA), a synthetic construct containing a T7 RNA polymerase promoter (ϕ2.5) followed by the mvRNA sequence was synthesized (Thermo Scientific GeneArt). The construct was flanked by AdeI and XbaI restriction sites and cloned into AdeI/XbaI-digested pJET_Gluc1 plasmid (29).

To generate plasmids encoding sense and antisense strands of dsRNA of defined lengths, fragments corresponding to portions of the Gaussia luciferase coding sequence (100-500 bp) or a fragment of the pKK-TEV-EGFP plasmid (1600 bp) (51) were amplified by PCR using pJET_Gluc1 or pJET_Gluc2 (for sense and antisense strands, respectively), or pKK-TEV-EGFP plasmid as templates, respectively. Amplification was performed using Phusion™ High-Fidelity DNA Polymerase (Thermo Scientific F530) and primers listed in Table S1.

PCR products were cloned into pJET1.2 using the CloneJET PCR Cloning Kit (Thermo Scientific K1231). Subsequently, pJET1.2 plasmids containing inserts encoding sense or antisense RNA strands were digested with AdeI (Thermo Scientific FD1234) and XbaI (Thermo Scientific FD0685). The inserts were then ligated into AdeI/XbaI-digested pJET_Gluc1 using T4 DNA ligase (Thermo Scientific EL0013), generating pJET_dsRNA_1 and pJET_dsRNA_2 constructs encoding sense and antisense strands of dsRNA of defined lengths, respectively.

The pSpCas9(BB)-2A-Puro (PX459) V2.0 plasmid (gift from Feng Zhang; Addgene plasmid #62988 (52)) was used to generate MDA5 knockout human and chicken cell lines. Overlapping oligonucleotides (Table S1) were annealed in T4 DNA ligase buffer and ligated into BpiI-digested pSpCas9(BB)-2A-Puro (PX459) V2.0 using T4 DNA ligase, generating three independent pSpCas9(BB)-2A-Puro-hMDA5 and three pSpCas9(BB)-2A-Puro-chMDA5 constructs.

### Generation of MDA5 knockout cell lines

A549 and DF-1 cells were seeded in 6-well plates and cultured to approximately 80% confluency prior to transfection. Cells were transfected with an equimolar mixture of three pSpCas9(BB)-2A-Puro-hMDA5 or pSpCas9(BB)-2A-Puro-chMDA5 plasmids (2 µg total DNA per well) using Lipofectamine 3000 (Invitrogen, L3000001).

In A549 cells, the culture medium was replaced 48 h post-transfection with fresh medium containing puromycin (1.5 µg/mL; InvivoGen ant-pr-1) for selection. After 3 days of selection, cells were cultured in antibiotic-free medium.

In DF-1 cells, the culture medium was replaced 24 h post-transfection with fresh medium containing puromycin (1 µg/mL) for selection. After 5 days of selection, cells were cultured in antibiotic-free medium. Once cells reached approximately 80% confluency, they were serially diluted to obtain single-cell clones in 96-well plates. Individual clones were expanded and screened for loss of MDA5 expression. In A549 cells, knockout efficiency was assessed by western blotting using anti-MDA5 antibodies. In DF-1 cells, due to the lack of suitable antibodies recognizing chicken MDA5, knockout clones were identified functionally by assessing the absence of immune activation following dsRNA stimulation, as measured by MX1 protein expression.

### T7 endonuclease I assay

Genomic DNA from monoclonal DF-1 MDA5 knockout (KO) lines and wild-type (WT) cells was isolated using the Cell Culture DNA Purification Kit (EURx E3555) according to the manufacturer’s instructions. The genomic region encompassing the target site was amplified by PCR using specific primers (Table S1) and Phusion™ High-Fidelity DNA Polymerase (Thermo Scientific, F530L). PCR products were purified using the PCR/DNA Clean-Up Purification Kit (EURx E3520) according to the manufacturer’s instructions.

Purified PCR products, either individually or as a 1:1 equimolar mixture of amplicons derived from WT and KO genomic DNA, were subjected to denaturation and reannealing to allow heteroduplex formation. Samples were then treated with T7 endonuclease I (2 U; EURx E1125) for 15 min at 37°C. Reaction products were resolved in an agarose gel in 1x TBE buffer.

### Amplicon deep sequencing

PCR amplicons were fragmented to an average size of ∼500 bp using a Covaris S220 instrument. Sequencing libraries were prepared from 350 ng of fragmented DNA using the KAPA Hyper Prep Kit (Roche KK8504) and KAPA UDI Indexes (KAPA Biosystems 08098140702) according to the manufacturer’s instructions. Libraries were amplified by seven cycles of PCR and size-selected to approximately 600 bp using KAPA Pure Beads (Roche 07983298001). Library quality and fragment size distribution were assessed using the Agilent TapeStation 4150, and library concentrations were determined by qPCR using the KAPA Library Quantification Kit (Kapa Biosciences KK4824). Equimolar pools of libraries were sequenced on the Illumina NovaSeq 6000 platform in paired-end mode (2 x 100 bp). Adapter trimming and quality filtering were performed using Cutadapt (v2.6) and Trimmomatic (v0.4), respectively. Reads were mapped using BWA-MEM (v0.7.19). Mapped reads were visualized using Integrative Genomics Viewer (IGV).

### In vitro transcription

RNA used in this study (Table S2) was generated by in vitro transcription as described previously (29), with modifications. Plasmid templates based on pJET constructs were linearized with PaqCI (New England Biolabs R0745) and used for transcription.

In vitro transcription reactions were performed in RNA polymerase buffer containing 40 mM Tris-HCl (pH 7.9), 10 mM MgCl_2_, 1 mM DTT, and 2 mM spermidine. Each reaction contained 40 ng/μL linearized DNA template, 2 mM each of ATP, CTP, GTP, and UTP, 1 U/μL RiboLock RNase inhibitor (Thermo Scientific EO0382), in-house-purified T7 RNA polymerase, and 1 U/μL pyrophosphatase (Thermo Scientific EF0221). Reactions were incubated at 37°C for 2 h, followed by treatment with DNase I (0.1 U/μL; Thermo Scientific EN0521) for 30 min at 37°C.

For generation of 50 bp dsRNA, a double-stranded DNA template was assembled by annealing overlapping oligonucleotides encoding the T7 RNA polymerase promoter (ϕ2.5) and a fragment of human leucine tRNA. The resulting DNA duplex was used directly as a template for in vitro transcription at a final concentration of 0.1 μM. All other reaction components and conditions were identical to those described above.

Crude RNA was purified using the Monarch® RNA Cleanup Kit (New England Biolabs, T2040) and further purified by high-performance liquid chromatography (HPLC) using an Agilent 1260 Infinity system equipped with an XBridge Premier Oligonucleotide BEH C18 column (130 Å, 2.5 μm, 4.6 x 150 mm; Waters 186009903).

HPLC purification was performed at 55°C using a linear gradient of buffer B (0.1 M triethylammonium acetate [TEAA], pH 7.0, and 50% acetonitrile) in buffer A (0.1 M TEAA, pH 7.0) over 30 min at a flow rate of 0.9 mL/min. Gradient conditions were adjusted depending on RNA length: 14.0-24.0% for mvRNA and 50 nucleotides (nt) RNA, 14.0-28.0% buffer B for 100-200 nt RNA, 21.9-26.5% buffer B for 250-550 nt RNA, and 24.0-28.6% for 1600 nt RNA. RNA from collected fractions was recovered by isopropanol precipitation.

To generate dephosphorylated transcripts, HPLC-purified RNA was treated with calf intestinal alkaline phosphatase (Invitrogen 18009019) at 2 U per 40 μL reaction (or 10 U per 40 μL for mvRNA) in the manufacturer’s buffer. Reactions were incubated for 1 h at 50°C (mvRNA) or for 30 min at 37°C (other RNAs), followed by purification using the Monarch® RNA Cleanup Kit.

### Double-stranded RNA (dsRNA) preparation

dsRNA was generated by annealing equimolar amounts of sense and antisense RNA strands. RNA strands (360 ng/μL each) were mixed in RNA annealing buffer (10 mM Tris-HCl, pH 7.0, 150 mM NaCl, 1 mM EDTA), heated to 90°C for 3 min, and then gradually cooled to room temperature.

Efficiency of duplex formation was assessed by agarose gel electrophoresis in 1x TBE buffer followed by ethidium bromide staining.

In cases where the distinction between single-stranded and double-stranded RNA was not clearly resolved by agarose gel electrophoresis, formation of dsRNA was further verified by RNase I digestion. Briefly, 250 ng of single-stranded or double-stranded RNA was incubated in a 10 μL reaction containing RNase I buffer (100 mM Tris-HCl, pH 7.5, 10 mM NaCl, 0.1 mM EDTA) and 0.1 U RNase I (Thermo Scientific, EN0601) for 5 min at 37°C. Reaction products were analyzed by agarose gel electrophoresis in 1x TBE buffer followed by ethidium bromide staining.

### RNA biotinylation

Biotinylated RNA was generated by in vitro transcription as described above, with the modification that CTP was partially substituted with Biotin-11-CTP (Lumiprobe 3354). Specifically, reactions contained 1.33 mM CTP and 0.67 mM Biotin-11-CTP instead of 2 mM CTP.

Crude RNA was purified using the Monarch® RNA Cleanup Kit and further purified by HPLC as described above, with modifications to the elution gradients for biotinylated transcripts. For biotinylated mvRNA, a linear gradient of buffer B (0.1 M triethylammonium acetate [TEAA], pH 7.0, and 50% acetonitrile) from 15% to 50% in buffer A (0.1 M TEAA, pH 7.0) over 30 min at a flow rate of 0.9 mL/min was used. For biotinylated 1600 nt RNA strands, a gradient of buffer B from 0% to 100% in buffer A over 30 min at a flow rate of 0.9 mL/min was applied. RNA from collected fractions was recovered by isopropanol precipitation.

Biotin incorporation was assessed by dot blot analysis. 1 μL of HPLC-purified RNA were spotted onto a Hybond-N+ membrane (Amersham 10320175). Following UV crosslinking, membranes were blocked overnight at 4°C with 3% BSA in PBST (PBS containing 0.1% Tween-20). Blots were incubated with streptavidin-HRP (Thermo Scientific 21130) diluted 1:6000 in PBST for 1 h at room temperature. Detection was performed using SuperSignal™ West Pico PLUS chemiluminescent substrate (Thermo Scientific 35065) and visualized using a ChemiDoc MP Imaging System (Bio-Rad). Following chemiluminescence detection, membranes were stained with methylene blue (0.02% in 0.3 M sodium acetate, pH 5.2).

To generate dephosphorylated transcripts, HPLC-purified RNA was treated with calf intestinal alkaline phosphatase at 2 U per 40 μL reaction for 1600 nt RNA or 10 U per 40 μL for mvRNA in the manufacturer’s buffer. Reactions were incubated for 30 min at 50°C (mvRNA) or 37°C (1600 nt RNA), followed by purification using the Monarch® RNA Cleanup Kit.

Biotinylated 1600 bp dsRNA was generated by annealing equimolar amounts of sense and antisense RNA strands. RNA strands (360 ng/μL each) were mixed in RNA annealing buffer, heated to 90°C for 3 min, and then gradually cooled to room temperature. Efficiency of duplex formation was assessed by agarose gel electrophoresis in 1x TBE buffer followed by ethidium bromide staining.

### Luciferase assay

A549, DF-1, and CCL-141 cells were seeded in 6-well plates and cultured to approximately 80% confluency prior to transfection. Cells were transfected with 1.5 μg of reporter plasmid using Lipofectamine 3000. For human cells, the IFN-β promoter-driven firefly luciferase reporter (IFN-Beta_pGL3; gift from Nicolas Manel, Addgene plasmid #102597 (53)) was used. For avian cells, a firefly luciferase reporter under the control of the chicken MX promoter (pMX (54)) was used.

Twenty-four hours post-transfection, cells were reseeded into 96-well plates. After an additional 24 h, cells were transfected with 350 ng/mL of the indicated dsRNA using 2 μL mRNA Boost Reagent and 2 μL TransIT-mRNA Reagent (components of the TransIT-mRNA Transfection Kit; Mirus MIR 2250) in 100 μL Opti-MEM (Gibco 51985026) per 1 mL of culture medium. Mock-transfected cells were treated identically but without dsRNA.

At 24 h post-dsRNA transfection, luciferase activity was measured using the Pierce™ Firefly Luc One-Step Glow Assay Kit (Thermo Fisher Scientific 16196) according to the manufacturer’s instructions. Luminescence was quantified using an Infinite 200PRO microplate reader (Tecan).

### Western blotting

A549, DF-1, and CCL-141 cells were seeded in 24-well plates and cultured to approximately 80% confluency prior to transfection. Cells were transfected with indicated dsRNA as described above or treated with sodium arsenite for 30 min. At the indicated time points, cells were lysed in RIPA buffer (50 mM Tris-HCl, pH 8.0, 5 mM EDTA, 1% Igepal CA-630, 0.5% sodium deoxycholate, 0.1% sodium dodecyl sulfate, 150 mM NaCl) or in Luciferase Cell Culture Lysis Reagent (Promega E1531) supplemented with cOmplete™ EDTA-free protease inhibitor cocktail (Roche 4693159001) and phosphatase inhibitor (Roche 4906845001). The latter was used for analysis of PKR and eIF2α phosphorylation.

Cell lysates were mixed with Laemmli sample buffer and heated at 95°C for 5 min. Proteins were separated by 10% SDS-polyacrylamide gel electrophoresis (SDS-PAGE) and transferred onto nitrocellulose membranes using the Mini Trans-Blot® Cell and Criterion™ Blotter (Bio-Rad).

Membranes were stained with Ponceau S to confirm equal loading and transfer, followed by blocking for 1 h in either 5% (w/v) skim milk in PBST or 5% (w/v) bovine serum albumin (BSA) in TBST. Membranes were incubated with primary antibodies diluted in PBST or TBST overnight at 4°C. After washing, membranes were incubated with HRP-conjugated secondary antibodies for 1 h at room temperature.

Signal detection was performed using SuperSignal™ West Pico PLUS chemiluminescent substrate and visualized with a ChemiDoc MP Imaging System.

### Puromycin incorporation assay

Puromycin incorporation assays were performed as described previously (55). Cells were seeded in 24-well plates and cultured to approximately 80% confluency prior to transfection with indicated dsRNA as described above or treated with sodium arsenite for 30 min. At 24 h post-transfection, puromycin was added to a final concentration of 4 μg/mL for 30 min prior to cell lysis in RIPA buffer. Lysates were prepared and analyzed by Western blotting as described above.

### RNase L activity analysis

A549, DF-1, and CCL-141 cells were seeded in 12-well plates and cultured to approximately 80% confluency prior to transfection. Cells were transfected with indicated dsRNA as described above.

Total RNA was isolated 24 h post-transfection using TRI Reagent™ Solution (Invitrogen AM9738), followed by purification with the Monarch® RNA Cleanup Kit.

RNA integrity and rRNA degradation were assessed either by electrophoresis in an agarose/0.4 M formaldehyde gel in NBC buffer (50 mM boric acid, 1 mM sodium acetate, 5 mM NaOH) followed by ethidium bromide staining, or using an Agilent 2100 Bioanalyzer with the RNA 6000 Nano Kit (Agilent 5067-1511).

### Virus infection

A total of 2 ×10^6^ A549 cells were infected with highly pathogenic avian influenza A virus H5N8 or the human-origin H1N1 virus at a multiplicity of infection (MOI) of 1 TCID50/cell (50% tissue culture infectious dose per cell). Infections were carried out in Opti-MEM medium supplemented with 1% penicillin-streptomycin and 1 µg/mL TPCK-treated trypsin (Thermo Scientific 20233). Cells were harvested when cytopathic effect reached approximately 50% (corresponding to 48 h post-infection for H1N1 and 72 h post-infection for H5N8). Cells were resuspended in 1 mL of TRIzol Reagent (Thermo Scientific 15596026) and stored at -70°C until RNA extraction.

### RNA fractionation

Total RNA isolated from virus-infected, dsRNA-treated, and mock-treated cells was subjected to HPLC fractionation using an Agilent 1260 Infinity system equipped with an XBridge Premier Oligonucleotide BEH C18 column.

Fractionation was performed at 55°C using a linear gradient of buffer B (0.1 M triethylammonium acetate [TEAA], pH 7.0, and 50% acetonitrile) in buffer A (0.1 M TEAA, pH 7.0), ranging from 20% to 29% buffer B over 30 min at a flow rate of 0.9 mL/min.

The fraction corresponding to short RNA species was collected between 2.5 and 12 min. Collected fractions were concentrated using Amicon® Ultra centrifugal filters (3 kDa MWCO; Millipore UFC800308) to a final volume of approximately 0.5 mL, followed by isopropanol precipitation. For downstream transfections of human and chicken cells, 350 ng of the short RNA fraction per 1 mL of culture medium was used.

To generate dephosphorylated short RNA species, the HPLC-purified RNA fraction was treated with calf intestinal alkaline phosphatase at 10 U per 40 μL reaction in the manufacturer’s buffer. Reactions were incubated for 1 h at 50°C, followed by purification using the Monarch® RNA Cleanup Kit.

### Northern blot hybridization

Total RNA (1 μg) and short RNA fractions (140 ng) were separated in 12% polyacrylamide gels containing 8 M urea in 1x TBE buffer and transferred to a Hybond-N+ membrane by electrotransfer in 0.5x TBE.RNA was crosslinked to the membrane by UV irradiation. Membranes were hybridized with γ-^32^P 5’ end-labeled oligonucleotide probe in PerfectHyb Plus hybridization buffer (Merck H7033) at 42°C. Blots were processed according to standard procedures.

### Pull down analysis

A549, DF-1, and CCL-141 cells were seeded in 100 mm dishes and cultured to approximately 80% confluency prior to transfection. Cells were transfected as described above with an equimass mixture of 5’-ppp mvRNA and 5’-ppp 1600 bp dsRNA for 24 h. Mock-treated cells were included as controls.

Cells were washed with PBS and harvested by scraping in 1600 μL of lysis buffer per dish (20 mM Tris-HCl, pH 7.5, 150 mM NaCl, 2 mM MgCl_2_, 2 mM DTT, 0.2% IGEPAL CA-630, supplemented with cOmplete™ EDTA-free protease inhibitor cocktail). Lysates were homogenized by passing through a 26G needle seven times and clarified by centrifugation at 16000 x g for 10 min at 4°C.

Clarified lysate was divided into six aliquots (five 300 μL aliquots and one 100 μL aliquot). The 100 μL aliquot was reserved as an input control. The remaining aliquots were incubated with 200 ng of biotinylated dsRNA of defined lengths and/or 5’ end modifications (no RNA added to control samples) for 1 h at 4°C with rotation.

Pre-washed streptavidin magnetic beads (PerkinElmer CMG-227) were prepared by washing four times with lysis buffer lacking protease inhibitors. The pre-washed beads were then added to each sample (10 μL of 50% slurry per sample) and incubated for 30 min at 4°C with rotation. Following incubation, beads were washed four times with lysis buffer lacking protease inhibitors and resuspended in 50 μL of the same buffer. Bound proteins were eluted by treatment with RNase III (2 U; EURx E1340) for 30 min at 4°C.

### Mass spectrometry measurements and data analysis

Samples consisted of biotinylated RNA co-immunoprecipitation eluates provided in 50 µL of buffer containing 20 mM Tris-HCl (pH 7.5), 150 mM NaCl, 2 mM MgCl_2_, 2 mM DTT, and 0.2% IGEPAL CA-630. Proteins were denatured by adding reagents to final concentrations of 2% sodium deoxycholate (SDC), 100 mM Tris-HCl, 5 mM tris(2-carboxyethyl)phosphine (TCEP), and 10 mM 2-chloroacetamide (CAA), supplemented with protease and phosphatase inhibitors. Samples were sonicated for 10 min at room temperature and incubated at 60°C for 30 min with shaking at 600 rpm.

Protein concentration was not determined due to the limited amount of material; instead, sample input was assessed by SDS-PAGE followed by silver staining. Proteins were digested using a modified SP3 on-bead Lys-C/trypsin workflow. Peptides were dried in a SpeedVac without heating and stored at -80°C until LC-MS/MS analysis.

Approximately 0.1 µg of peptides was separated on a Waters C18 column (0.3 mm x 150 mm, 2.6 µm particle size) using a Micro M5 SCIEX LC system at a flow rate of 5 µL/min in direct-injection mode. Mass spectrometry analysis was performed on a ZenoTOF 7600 instrument (SCIEX) equipped with an OptiFlow Turbo V ion source and a microflow probe.

Source parameters were as follows: ionspray voltage 5000 V, source temperature 200°C, curtain gas 35 psi, gas 1 at 20 psi, and gas 2 at 60 psi. Data-dependent acquisition was performed using top 20-100 precursor selection with dynamic background subtraction. The TOF MS accumulation time was 100 ms, and MS/MS spectra were acquired at 30 V collision energy with a 5 ms accumulation time.

Raw LC-MS/MS data were processed using MaxQuant (v2.6.8.0) with the integrated Andromeda search engine. Because the experiment included human, chicken, and duck cell lines, database searches were performed separately for each species using UniProtKB FASTA files downloaded in May 2025: *Homo sapiens* (UP000005640) for A549 cells, *Gallus gallus* (UP000000539) for DF-1 cells, and *Anas platyrhynchos* (taxonomy ID 8839) for CCL-141 cells.

Label-free quantification (LFQ) was performed using razor peptides. Contaminants were included during database searching and removed prior to statistical analysis. Searches were conducted with a 1% false discovery rate (FDR) at both peptide-spectrum match (PSM) and protein levels, using sequence-reversed decoy databases. A minimum peptide length of seven amino acids was required, and the match-between-runs feature was enabled.

ProteinGroups outputs from MaxQuant were analyzed separately for each cell line using LFQ-Analyst in R. Proteins annotated as contaminants, reverse hits, or “only identified by site,” as well as proteins identified by a single peptide or not consistently quantified within a condition, were excluded. LFQ intensities were log2-transformed and normalized under the assumption that most proteins were unchanged. Missing values were imputed using a missing-not-at-random approach with a left-shifted Gaussian distribution (shift = 1.8 standard deviations, width = 0.3).

Pairwise differential protein abundance analyses were performed using limma empirical Bayes linear models. Proteins with a Benjamini-Hochberg adjusted p-value ≤ 0.05 and an absolute log2 fold change ≥ 1 were considered significantly regulated.

The mass spectrometry proteomic data have been deposited to the ProteomeXchange Consortium via the PRIDE partner repository with the dataset identifiers PXD077989, PXD078001, and PXD078031.

## Acknowledgments

The pSpCas9(BB)-2A-Puro (PX459) V2.0 plasmid was a gift from F. Zhang (Addgene plasmid no. 62988), the IFN-Beta_pGL3 plasmid was a gift from N. Manel (Addgene plasmid no. 102597), and the reverse-genetics plasmids for influenza virus A/WSN/1933 (H1N1) were a gift from Gülsah Gabriel.

This work was supported by the National Science Centre (grant nos. 2021/42/E/NZ1/00314 and 2025/57/B/NZ9/02722) and the National Research and Development Centre (ICRAD/II/76/FLU-SWITCH/2023) awarded to P.J.S., and ICRAD, an ERA-NET co-funded under European Union’s Horizon 2020 research and innovation programme, under Grant Agreement no. 862605 to R.V. and P.J.S. Next-generation sequencing was performed at the Genomics Core Facility CeNT UW (RRID:SCR_022718) using the NovaSeq 6000 platform, financed by the Polish Ministry of Science and Higher Education (decision no. 6817/IA/SP/2018, dated 10 April 2018).

