## Supporting Information for "RNA length and receptor usage define innate immune recognition across species"

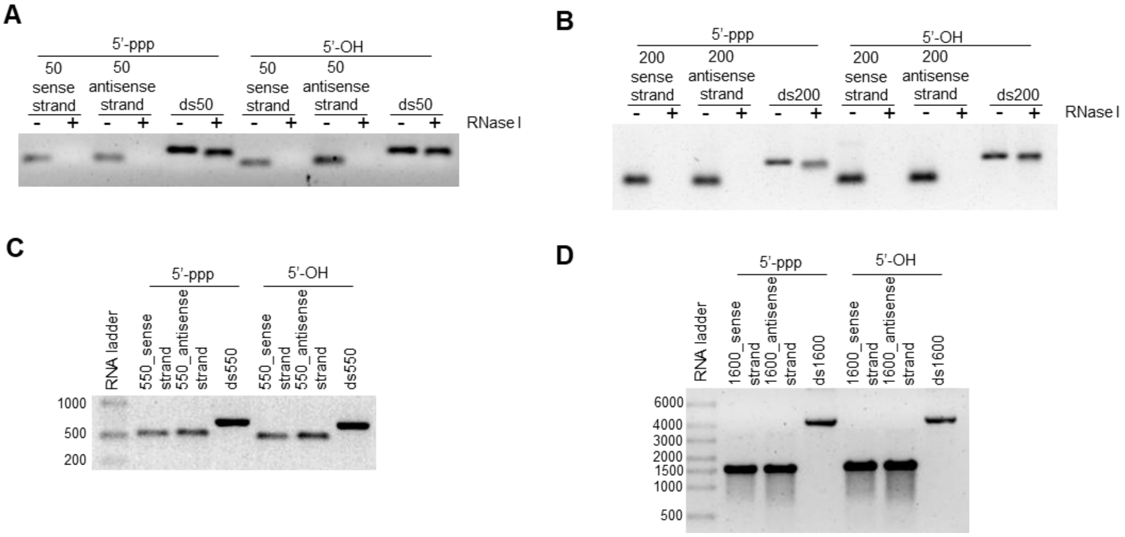


**Fig. S1. Preparation of dsRNA molecules.** (A) Stability analysis of 50 nt sense and antisense RNA strands bearing either a 5'-triphosphate (5'-ppp) or a dephosphorylated (5'-OH) end, and duplexes formed from these strands. Transcripts were incubated with or without RNase I, and products were analyzed in a native agarose gel. (B) Stability analysis of 200 nt sense and antisense RNA strands bearing either a 5'-ppp or 5'-OH end, and corresponding duplexes, following RNase I treatment as in (A). (C) Analysis of 550 nt sense and antisense RNA strands bearing either a 5'-ppp or 5'-OH end, and duplexes formed from these strands, resolved in a native agarose gel. (D) Analysis of 1600 nt sense and antisense RNA strands bearing either a 5'-ppp or 5'-OH end, and duplexes formed from these strands, resolved in a native agarose gel.

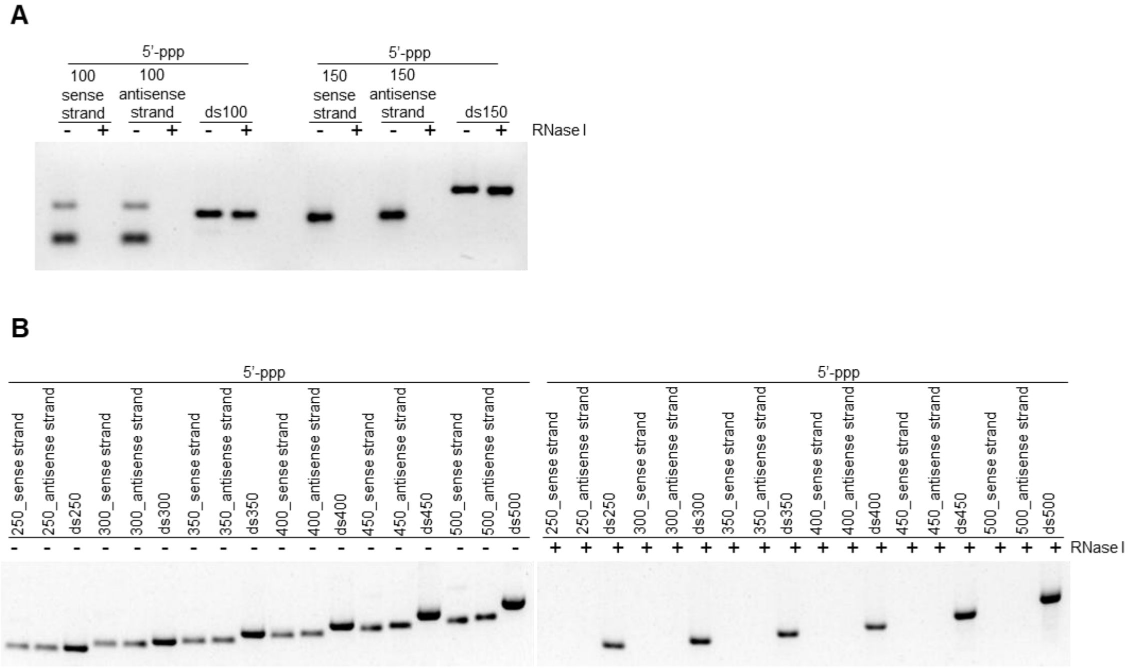


**Fig. S2. Preparation of dsRNA molecules.** (A) Stability analysis of 100 nt and 150 nt sense and antisense RNA strands bearing either a 5'-triphosphate (5'-ppp) end, and duplexes formed from these strands. Transcripts were incubated with or without RNase I, and products were analyzed in a native agarose gel. (B) Stability analysis of 250 nt, 300 nt, 350 nt, 400 nt, 450 and 500 nt sense and antisense RNA strands bearing either a 5'-ppp or 5'-OH end, and corresponding duplexes, following RNase I treatment as in (A).

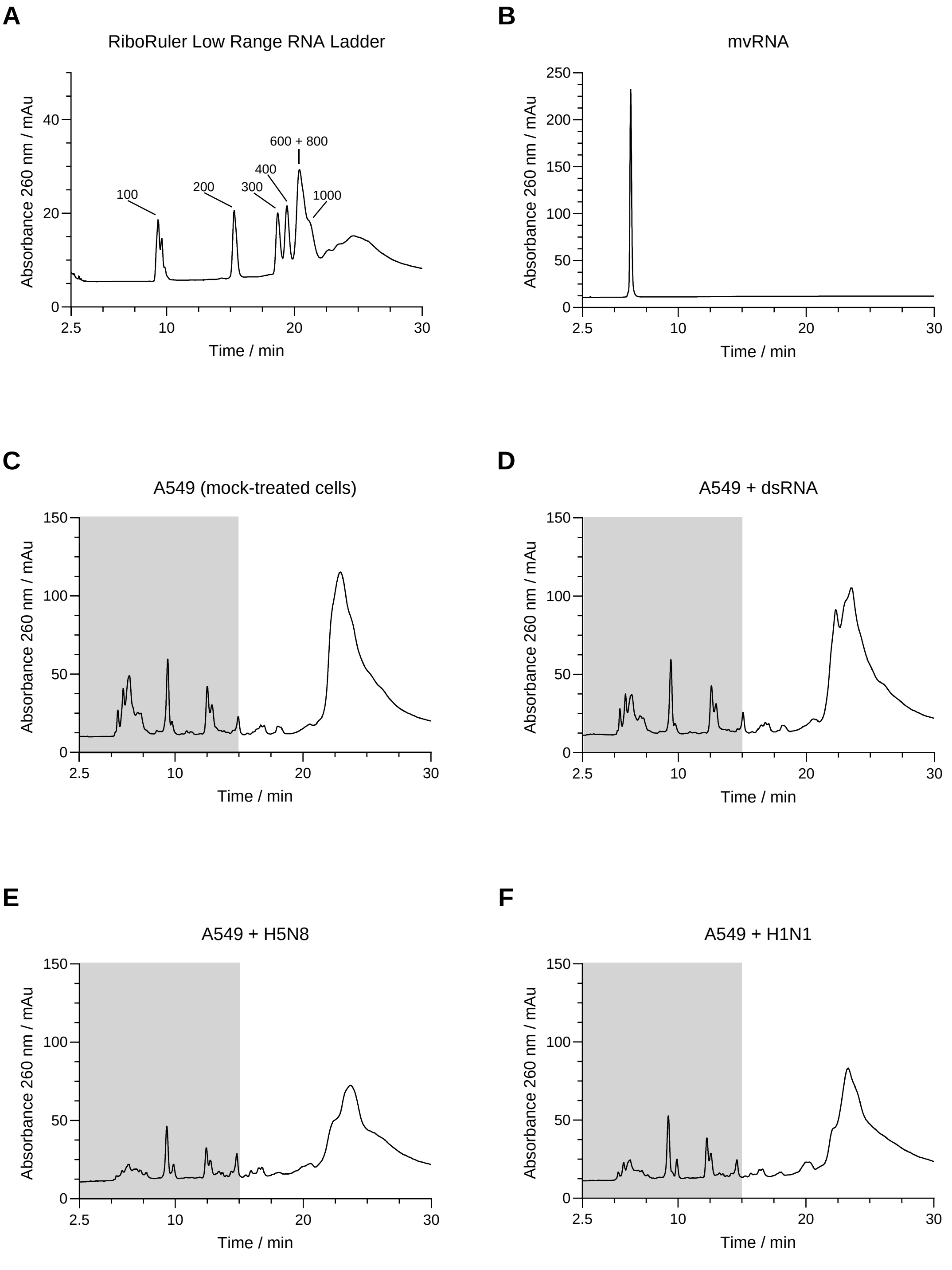


**Fig. S3. HPLC fractionation.** (A) Representative chromatogram of the RiboRuler Low Range RNA Ladder (Thermo Scientific, SM1831). RNA species of defined lengths were assigned based on their retention times. (B) Representative HPLC chromatogram of an in vitro transcribed mvRNA (60 nt). (C-F) Fractionation profiles of total RNA isolated from mock-treated, dsRNA-treated, and virus

(H5N8 or H1N1)-infected A549 cells. The collected fraction corresponding to short RNA species is highlighted in gray.

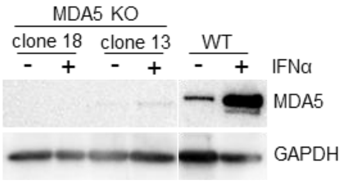


**Fig. S4. Verification of MDA5 knockout in A549 cells.** Wild-type (WT) and MDA5 knockout (KO) A549 cells were treated with IFNα (200 U/mL; Pestka Biomedical Laboratories 11200) for 24 h, and MDA5 expression was assessed by western blot analysis. KO clone 18 was used for subsequent experiments.

A

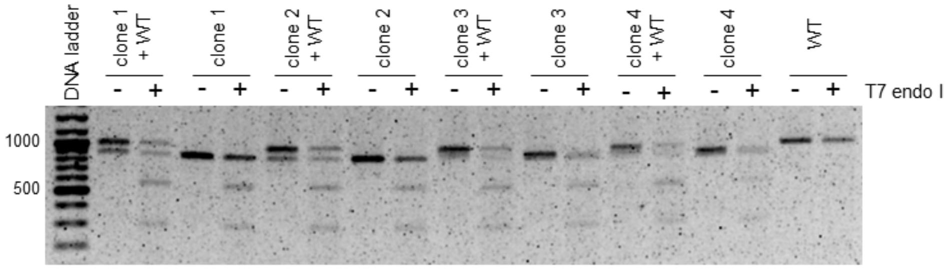
DNA ladder
clone 1 + WT
clone 1
clone 2 + WT
clone 2
clone 3 + WT
clone 3
clone 4 + WT
clone 4
WT
- + - + - + - + - + - + - + - + - +
T7 endo I
1000
500

B

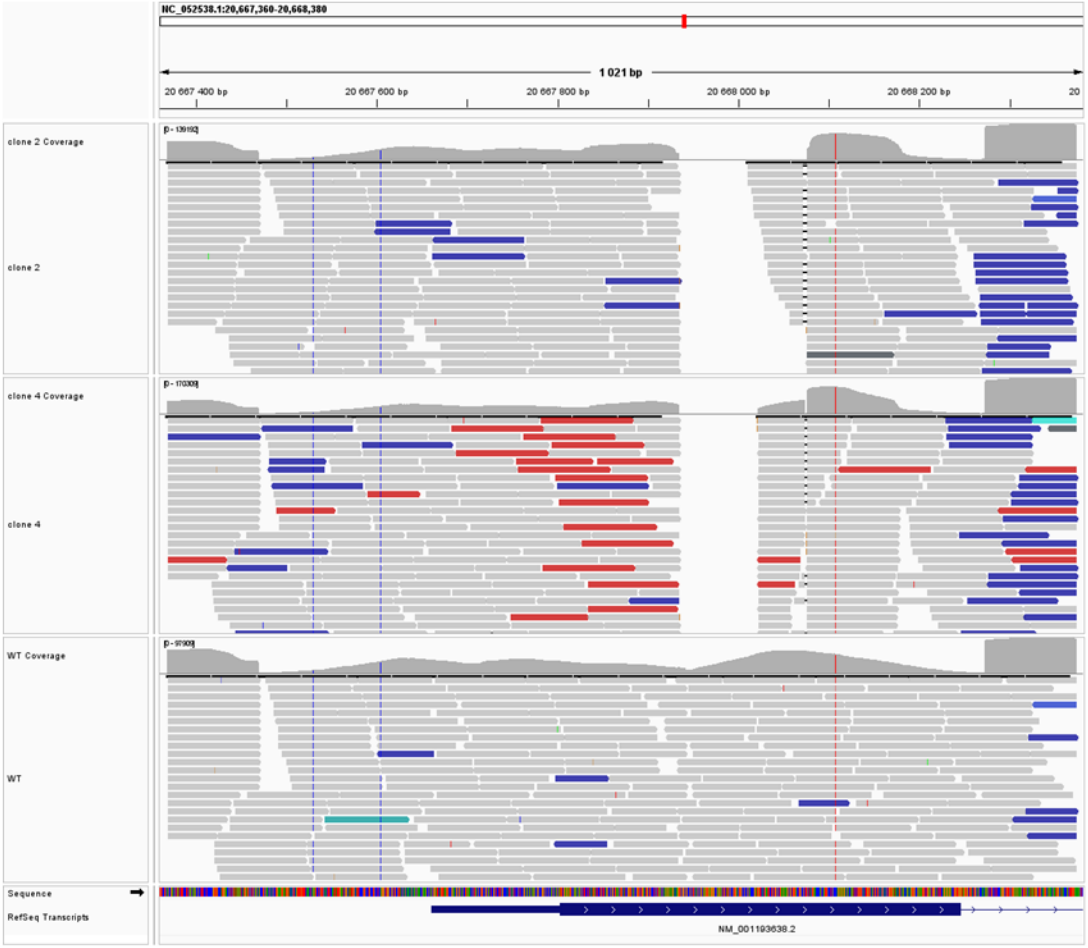
1 021 bp
clone 2 Coverage
clone 2
clone 4 Coverage
clone 4
WT Coverage
WT
Sequence
RefSeq Transcripts

C

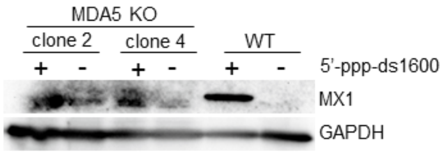
MDA5 KO
clone 2
clone 4
WT
+ - + - + -
5'-ppp-ds1600
MX1
GAPDH

**Fig. S5. Verification of MDA5 knockout in DF-1 cells.** (A) T7 endonuclease I (T7 endo I) assay validating CRISPR/Cas9-mediated genome editing. Purified PCR products, examined either individually or as a 1:1 equimolar mixture of amplicons derived from wiled-type (WT) and KO genomic DNA, were subjected to denaturation and reannealing to allow heteroduplex formation, followed by treatment with T7 endonuclease I. Reaction products were resolved in an agarose gel. (B) Representative IGV visualization of sequencing reads across the edited locus. Alignments are shown for the indicated samples. Coverage tracks are displayed above each alignment panel. (C) WT and MDA5 KO DF-1 cells were treated with 5'-ppp-ds550 for 24 h. Due to the lack of suitable antibodies recognizing chicken MDA5, functional validation of MDA5 deficiency was assessed by western blot analysis of MX1 expression following dsRNA stimulation. Clone 4 was used for subsequent experiments.

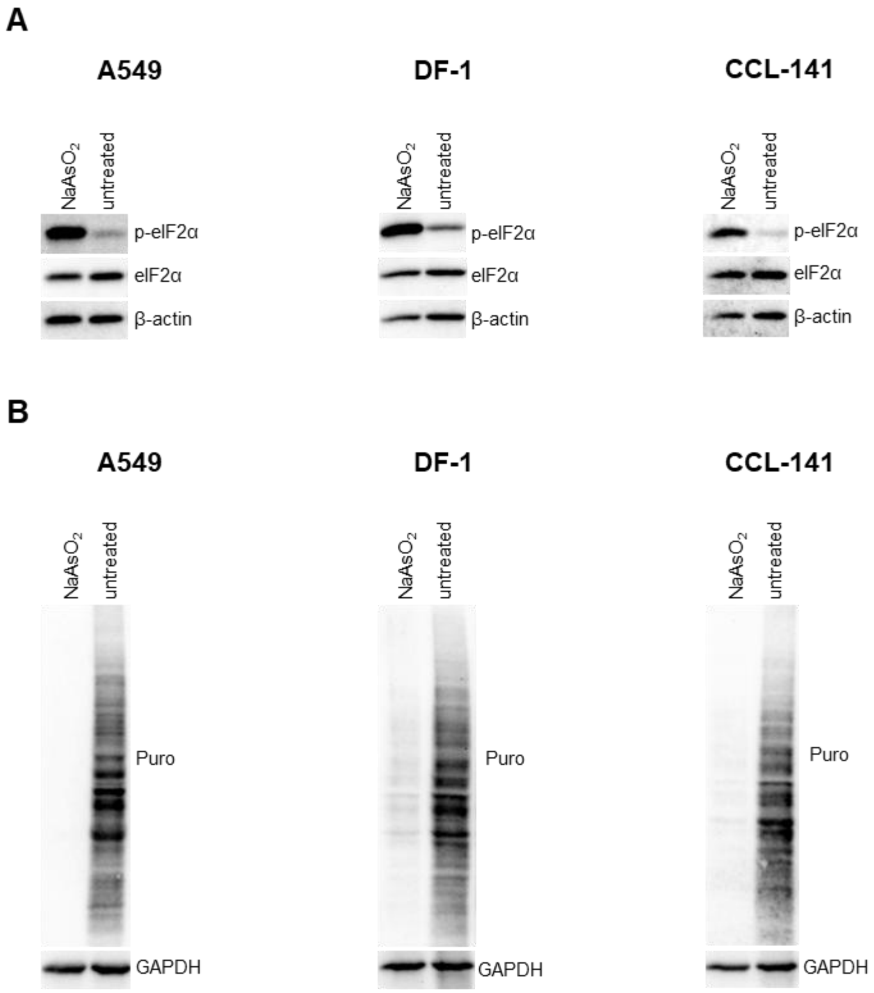


**Fig. S6. Validation of translational inhibition assays across species.** (A) Western blot analysis of eIF2α phosphorylation in human (A549), chicken (DF-1), and duck (CCL-141) cells following treatment with sodium arsenite ($NaAsO_2$). (B) Puromycin incorporation assay to assess global translation in the indicated cell lines under the same conditions. Reduced puromycin signal indicates inhibition of protein synthesis.

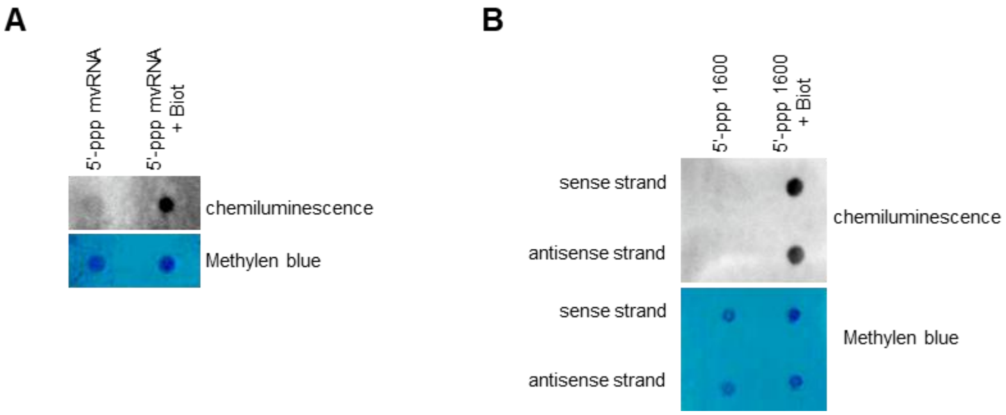


**Fig. S7. Validation of biotin incorporation during in vitro transcription.** Dot blot analysis of biotin incorporation into in vitro transcribed (A) mvRNA (300 ng) and (B) sense and antisense strands of 1600 bp dsRNA (100 ng). Methylene blue staining served as a loading control.

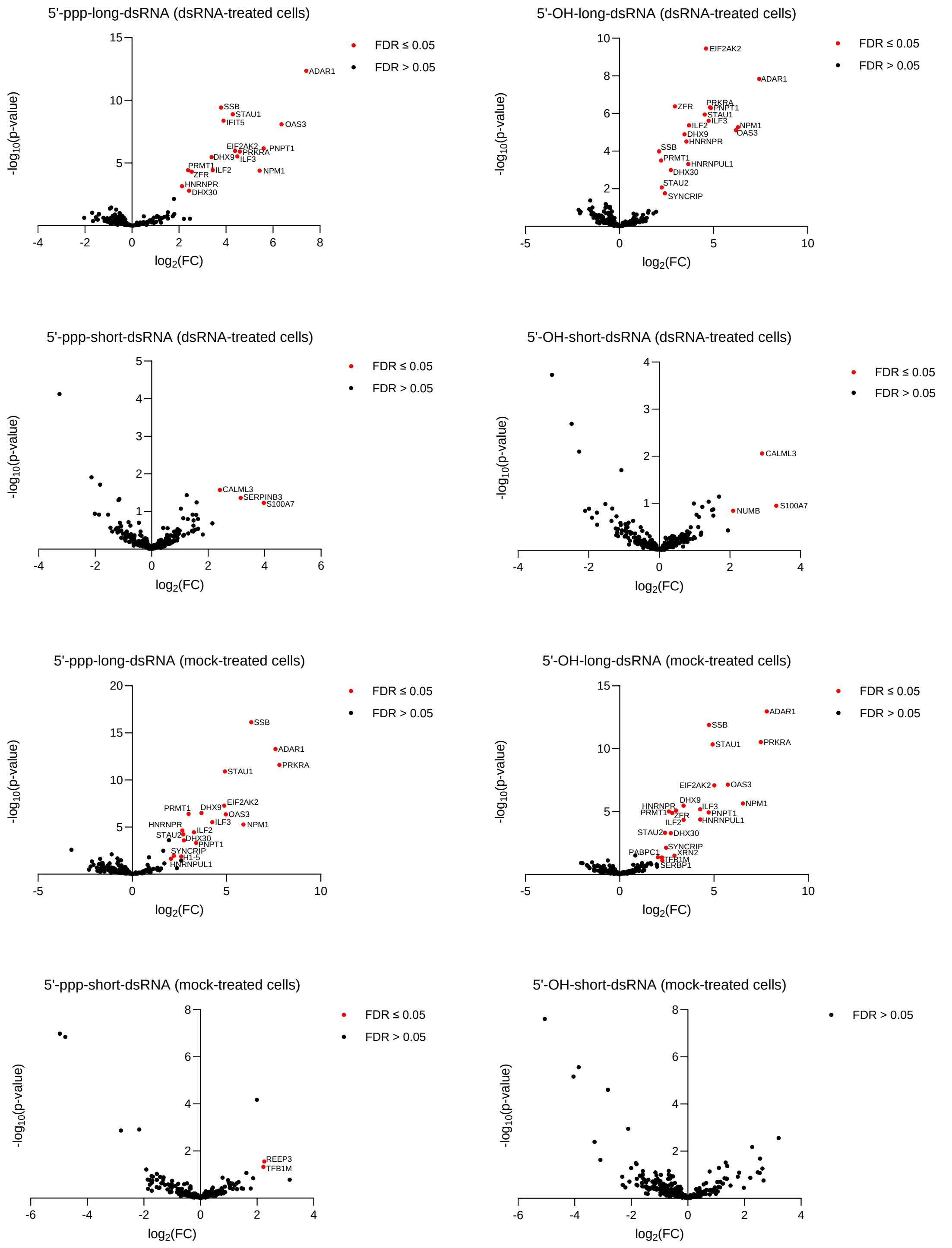


**Fig. S8. Volcano plots of proteins identified in pull-down analyses of A549 cell lysates.** The plots show log2(fold change) [log2(FC)] on the x-axis and −log10(p-value) on the y-axis. Proteins significantly enriched (FDR < 0.05) are highlighted in red and labeled, whereas rest of identified proteins is shown in black. For each plot, the dsRNA used as bait (short or long) and its 5' end modification (5’-ppp or 5’-OH), as well as the source of the lysate (mock-treated or dsRNA-treated cells) are indicated above the graph.

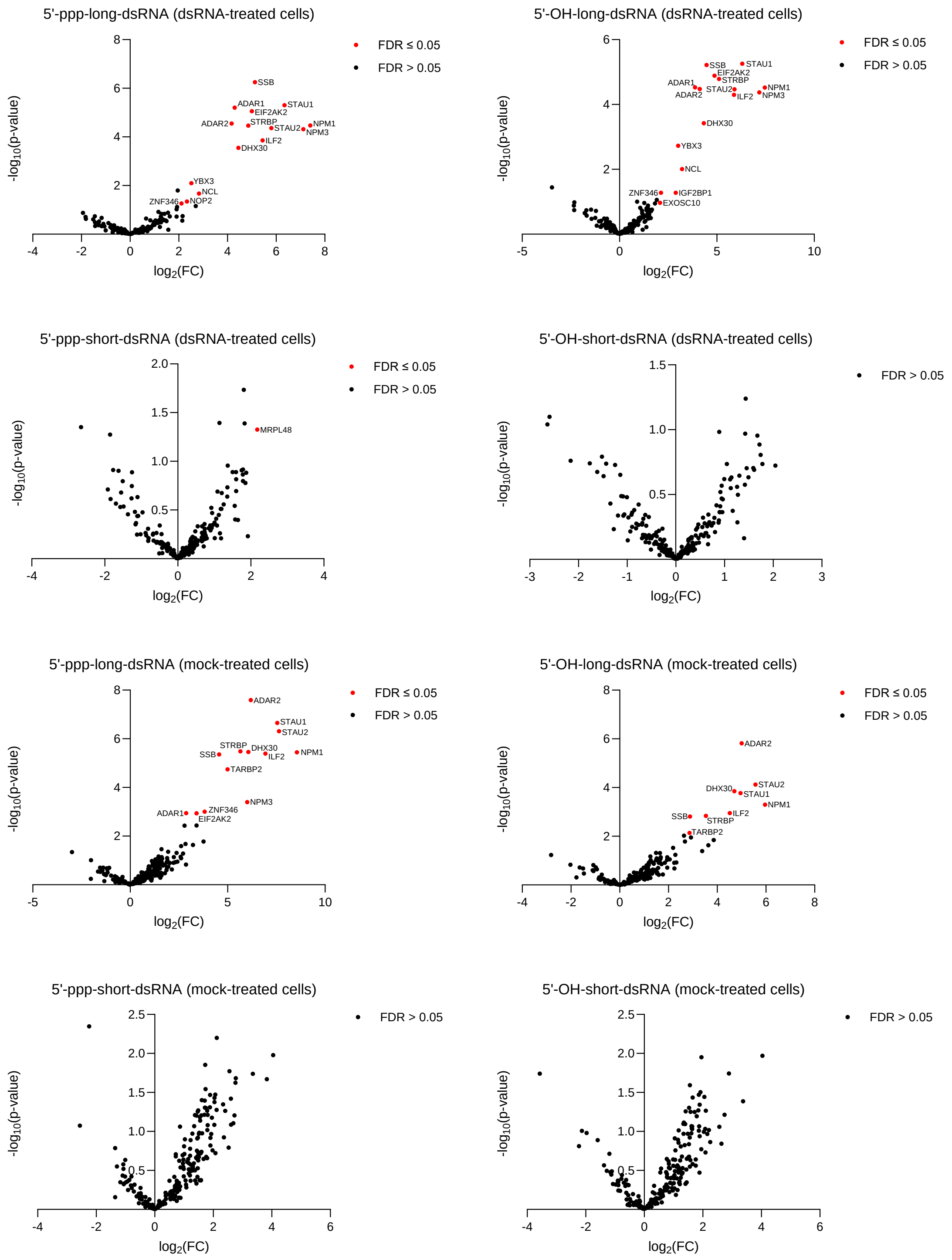


**Fig. S9. Volcano plots of proteins identified in pull-down analyses of DF-1 cell lysates.** The plots show log2(fold change) [log2(FC)] on the x-axis and −log10(p-value) on the y-axis. Proteins significantly enriched (FDR < 0.05) are highlighted in red and labeled, whereas rest of identified proteins is shown in black. For each plot, the dsRNA used as bait (short or long) and its 5’ end modification (5’-ppp or 5’-OH), as well as the source of the lysate (mock-treated or dsRNA-treated cells) are indicated above the graph.

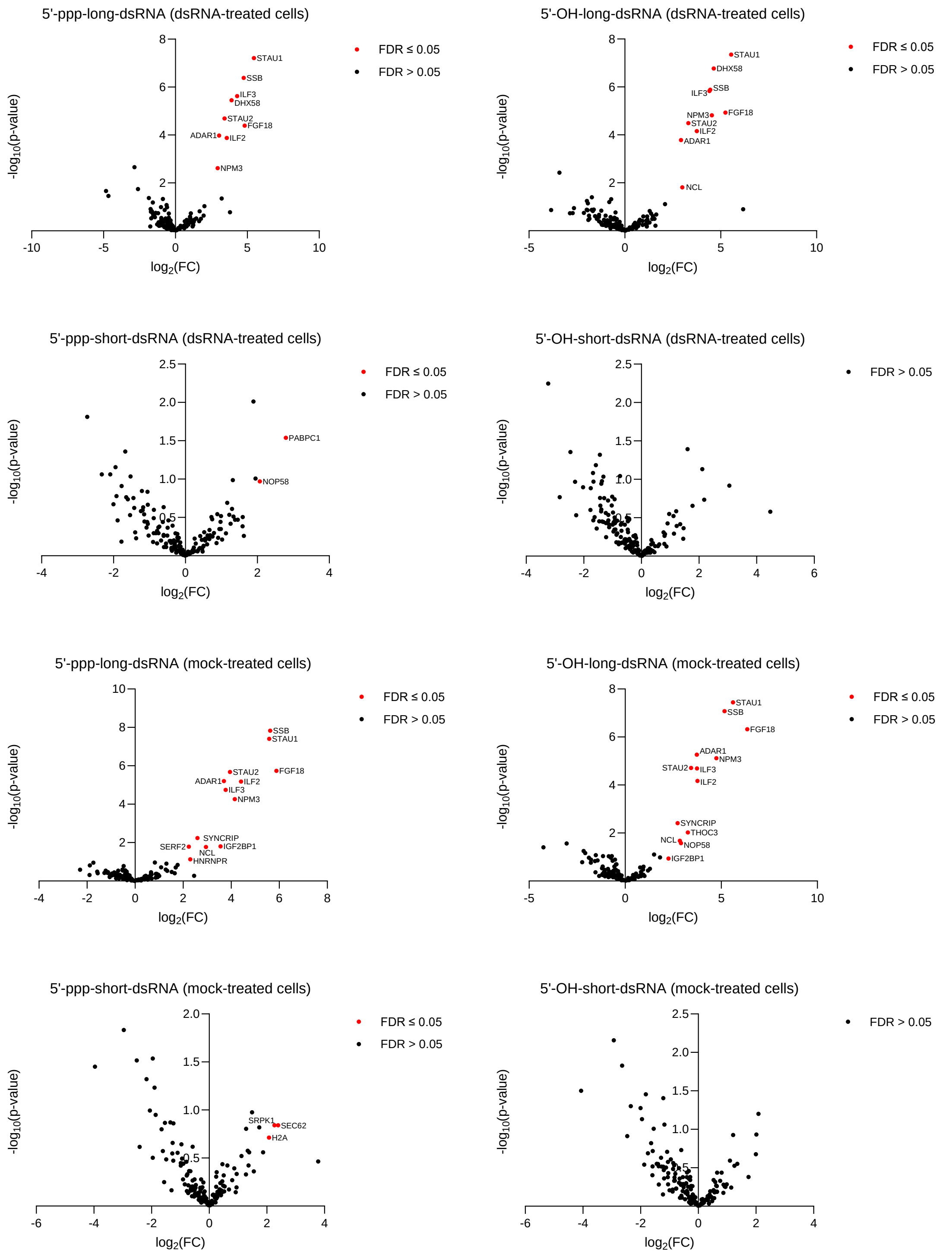


**Fig. S10. Volcano plots of proteins identified in pull-down analyses of CCL-141 cell lysates.** The plots show log2(fold change) [log2(FC)] on the x-axis and −log10(p-value) on the y-axis. Proteins significantly enriched (FDR < 0.05) are highlighted in red and labeled, whereas rest of identified proteins is shown in black. For each plot, the dsRNA used as bait (short or long) and its

5’ end modification (5’-ppp or 5’-OH), as well as the source of the lysate (mock-treated or dsRNA-treated cells) are indicated above the graph.

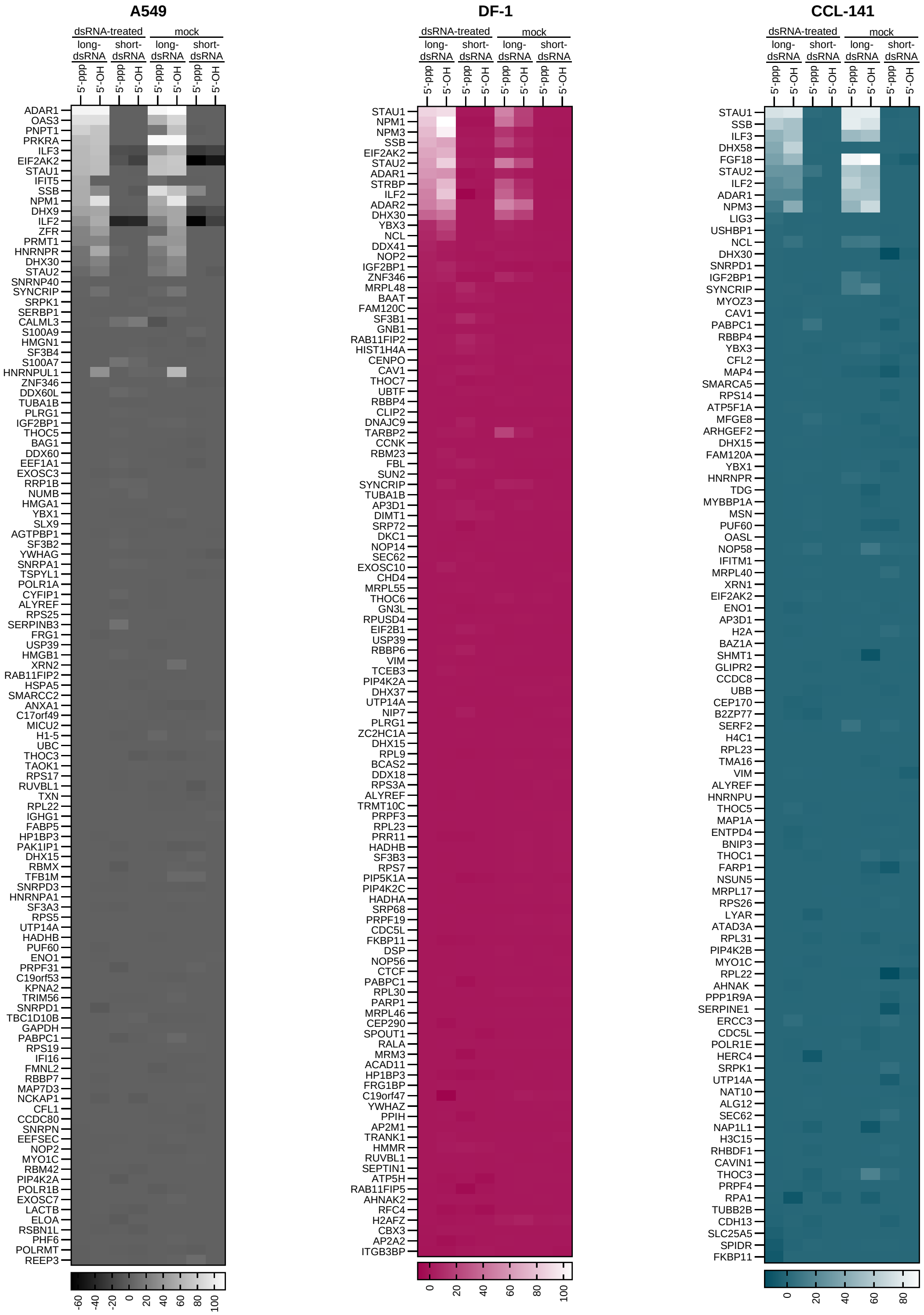


**Fig. S11. Full heatmaps corresponding to Fig. 6B.** Heatmaps showing enrichment of RNA-binding proteins in pull-down assays from human, chicken, and duck cell lysates. Protein enrichment is displayed as a composite score calculated as log2(FC) x -log10(p-value), relative to beads-only control samples. Color scales indicate relative protein enrichment. In contrast to Fig. 6B, which shows the top 30 proteins, this figure presents the complete dataset.

**Tables**

**Table S1. List of oligonucleotides used in this study.**

| Name | Sequence | Purpose |
|---|---|---|
| sgchMDA5_1f | CACCGTGAACCATCCGGGGTCGCGG | CRISPR/Cas9 gene knockout in A549 |
| sgchMDA5_1r | AAACCCGCGACCCCGGATGGTTCAC | CRISPR/Cas9 gene knockout in A549 |
| sgchMDA5_2f | CACCGGCTGGGGTTCACGTAGCAAG | CRISPR/Cas9 gene knockout in A549 |
| sgchMDA5_2r | AAACCTTGCTACGTGAACCCCAGCC | CRISPR/Cas9 gene knockout in A549 |
| sgchMDA5_3f | CACCGAGAGGAGAAGGACAAGGTGC | CRISPR/Cas9 gene knockout in A549 |
| sgchMDA5_3r | AAACGCACCTTGTCCTTCTCCTCTC | CRISPR/Cas9 gene knockout in A549 |
| sghMDA5_1f | CACCGCGTCATTGTCAGGCACAGAG | CRISPR/Cas9 gene knockout in DF-1 |
| sghMDA5_1r | AAACCTCTGTGCCTGACAATGACGC | CRISPR/Cas9 gene knockout in DF-1 |
| sghMDA5_2f | CACCGTGGTTGGACTCGGGAATTCG | CRISPR/Cas9 gene knockout in DF-1 |
| sghMDA5_2r | AAACCGAATTCCCGAGTCCAACCAC | CRISPR/Cas9 gene knockout in DF-1 |
| sghMDA5_3f | CACCGGGTGGAGCCTGTGCTGGACT | CRISPR/Cas9 gene knockout in DF-1 |
| sghMDA5_3r | AAACAGTCCAGCACAGGCTCCACCC | CRISPR/Cas9 gene knockout in DF-1 |
| ds250_sense_rev | GATCTTCTAGACACCTGCACGTAGGGT GCACTTGATGTGGGACAGGC | Cloning of sense strand of 250 bp dsRNA into pJET |
| ds250_antisense_for | CATCACGCTGTGTAATACGACTCACTAT TAGGGTGCACTTGATGTGGGACAGGC | Cloning of antisense strand of 250 bp dsRNA into pJET |
| ds300_sense_rev | GATCTTCTAGACACCTGCACGTAGGGG CCTTCGTAGGTGTGGCAGCG | Cloning of sense strand of 300 bp dsRNA into pJET |
| ds300_antisense_for | CATCACGCTGTGTAATACGACTCACTAT TAGGGGCCTTCGTAGGTGTGGCAGCG | Cloning of antisense strand of 300 bp dsRNA into pJET |
| ds350_sense_rev | GATCTTCTAGACACCTGCACGTAGGGG GAATGTCGACGATCGCCTC | Cloning of sense strand of 350 bp dsRNA into pJET |
| ds350_antisense_for | CATCACGCTGTGTAATACGACTCACTAT TAGGGGGAATGTCGACGATCGCCTC | Cloning of antisense strand of 350 bp dsRNA into pJET |
| ds400_sense_rev | GATCTTCTAGACACCTGCACGTAGGGG TGCGATGAACTGCTCCAAGGG | Cloning of sense strand of 400 bp dsRNA into pJET |
| ds400_antisense_for | CATCACGCTGTGTAATACGACTCACTAT TAGGGGTGCGATGAACTGCTCCAAGG | Cloning of antisense strand of 400 bp dsRNA into pJET |
| ds450_sense_rev | GATCTTCTAGACACCTGCACGTAGGGG GCAAGCCCTTTGAGGCAG | Cloning of sense strand of 450 bp dsRNA into pJET |
| ds450_antisense_for | CATCACGCTGTGTAATACGACTCACTAT TAGGGGGCAAGCCCTTTGAGGCA | Cloning of antisense strand of 450 bp dsRNA into pJET |
| ds500_sense_rev | GATCTTCTAGACACCTGCACGTAGGGG CACAGCGTTGCGGCAGCCAC | Cloning of sense strand of 500 bp dsRNA into pJET |
| ds500_antisense_for | CATCACGCTGTGTAATACGACTCACTAT TAGGGGCACAGCGTTGCGGCAGCCAC | Cloning of antisense strand of 500 bp dsRNA into pJET |
| ds100_sense_rev | GAT CTT CTA GAC ACC TGC ACG TAG GGT GGC CAC GGC CAC GAT GTT G | Cloning of sense strand of 100 bp dsRNA into pJET |

| | | |
|---|---|---|
| ds100_antisense_for | CATCACGCTGTGTAATACGACTCACTAT TAGGGTGGCCACGGCCACGATGTTG | Cloning of antisense strand of 100 bp dsRNA into pJET |
| ds150_sense_rev | GAT CTT CTA GAC ACC TGC ACG TAG GGG CCG GGC AAC TTC CCG CGG TC | Cloning of sense strand of 150 bp dsRNA into pJET |
| ds150_antisense_for | CATCACGCTGTGTAATACGACTCACTAT TAGGGGCCGGGCAACTTCCCGCGGTC | Cloning of antisense strand of 150 bp dsRNA into pJET |
| ds200_sense_rev | GAT CTT CTA GAC ACC TGC ACG TAG GGT TCC GGG CAT TGG CTT CCA AC | Cloning of sense strand of 200 bp dsRNA into pJET |
| ds200_antisense_for | CATCACGCTGTGTAATACGACTCACTAT TAGGGTTCCGGGCATTGGCTTCCAAC | Cloning of antisense strand of 200 bp dsRNA into pJET |
| pJET_up_for | GCTCTGAAGTCTTCTTCATT | Cloning of sense strand of dsRNA of chosen length into pJET |
| pJET_down_rev | TGCTTCCGGCTCGTATAATG | Cloning of antisense strand of dsRNA of chosen length into pJET |
| ds50_sense_f | CAGTAATACGACTCACTATTAGGGTCGT TAGCTCAGTTGGTAGAGCAGTTGACTTT TAATCAATTGCCCT | Upon annealing with ds50_sense_r serves as a template for in vitro transcription |
| ds50_antisense_f | CAGTAATACGACTCACTATTAGGGCAAT TGATTAAAAGTCAACTGCTCTACCAACT GAGCTAACGACCCT | Upon annealing with ds50_antisense_r serves as a template for in vitro transcription |
| ds50_sense_r | AGGGCAATTGATTAAAAGTCAACTGCTC TACCAACTGAGCTAACGACCCTAATAGT GAGTCGTATTACTG | Upon annealing with ds50_sense_f serves as a template for in vitro transcription |
| ds50_antisense_r | AGGGTCGTTAGCTCAGTTGGTAGAGCA GTTGACTTTTAATCAATTGCCCTAATAG TGAGTCGTATTACTG | Upon annealing with ds50_antisense_f serves as a template for in vitro transcription |
| ds1600_sense_f | GCCACGCTGTGTAATACGACTCACTATT AGGGCCGAGCGCAGAAGTGG | Cloning of sense strand of 1600 bp dsRNA into pJET |
| ds1600_sense_r | GCTCTAGACACCTGCACGTAGGGAGAG CTCTGCTTATATAG | Cloning of sense strand of 1600 bp dsRNA into pJET |
| ds1600_antisense_f | CGTCTAGACACCTGCACGTAGGGCCGA GCGCAGAAGTGG | Cloning of antisense strand of 1600 bp dsRNA into pJET |
| ds1600_antisense_r | CGCACGCTGTGTAATACGACTCACTATT AGGGAGAGCTCTGCTTATATAG | Cloning of antisense strand of 1600 bp dsRNA into pJET |
| SNORD44 | AGTTAGAGCTAATTAAGACCT | Probe for northern blot hybridization |
| chMDA5_T7f | AGCCGGACGCTGATCTTTCC | PCR to verify MDA5 knockout in DF-1 cells by T7 endonuclease assay |
| chMDA5_T7r | TTCACAAAGCTGGGTTATGC | PCR to verify MDA5 knockout in DF-1 cells by T7 endonuclease assay |

**Table S2. RNA sequences used in this study.** For duplexes, only the sense strand sequence is shown.

| Name | Sequence |
|---|---|
| mvRNA | AGUAGAAACAAGGUCGUUUUUAAACUAUUCAACACUGAAUAUAAUUGACCUGC UUUCGCU |
| dsRNA 50 bp | AGGGUCGUUAGCUCAGUUGGUAGAGCAGUUGACUUUUAAUCAAUUGCCCU |
| dsRNA 100 bp | AGGGAGUCAAAGUUCUGUUUGCCCUGAUCUGCAUCGCUGUGGCCGAGGCCAA GCCCACCGAGAACAACGAAGACUUCAACAUCGUGGCCGUGGCCACCCU |
| dsRNA 150 bp | AGGGAGUCAAAGUUCUGUUUGCCCUGAUCUGCAUCGCUGUGGCCGAGGCCAA GCCCACCGAGAACAACGAAGACUUCAACAUCGUGGCCGUGGCCAGCAACUUC GCGACCACGGAUCUCGAUGCUGACCGCGGGAAGUUGCCCGGCCCCU |
| dsRNA 200 bp | AGGGAGUCAAAGUUCUGUUUGCCCUGAUCUGCAUCGCUGUGGCCGAGGCCAA GCCCACCGAGAACAACGAAGACUUCAACAUCGUGGCCGUGGCCAGCAACUUC GCGACCACGGAUCUCGAUGCUGACCGCGGGAAGUUGCCCGGCAAGAAGCUG CCGCUGGAGGUGCUCAAAGAGUUGGAAGCCAAUGCCCGGAACCCU |
| dsRNA 250 bp | AGGGAGUCAAAGUUCUGUUUGCCCUGAUCUGCAUCGCUGUGGCCGAGGCCAA GCCCACCGAGAACAACGAAGACUUCAACAUCGUGGCCGUGGCCAGCAACUUC GCGACCACGGAUCUCGAUGCUGACCGCGGGAAGUUGCCCGGCAAGAAGCUG CCGCUGGAGGUGCUCAAAGAGUUGGAAGCCAAUGCCCGGAAAGCUGGCUGCA CCAGGGGCUGUCUGAUCUGCCUGUCCCACAUCAAGUGCACCCU |
| dsRNA 300 bp | AGGGAGUCAAAGUUCUGUUUGCCCUGAUCUGCAUCGCUGUGGCCGAGGCCAA GCCCACCGAGAACAACGAAGACUUCAACAUCGUGGCCGUGGCCAGCAACUUC GCGACCACGGAUCUCGAUGCUGACCGCGGGAAGUUGCCCGGCAAGAAGCUG CCGCUGGAGGUGCUCAAAGAGUUGGAAGCCAAUGCCCGGAAAGCUGGCUGCA CCAGGGGCUGUCUGAUCUGCCUGUCCCACAUCAAGUGCACGCCCAAGAUGAA GAAGUUCAUCCCAGGACGCUGCCACACCUACGAAGGCCCCU |
| dsRNA 350 bp | AGGGAGUCAAAGUUCUGUUUGCCCUGAUCUGCAUCGCUGUGGCCGAGGCCAA GCCCACCGAGAACAACGAAGACUUCAACAUCGUGGCCGUGGCCAGCAACUUC GCGACCACGGAUCUCGAUGCUGACCGCGGGAAGUUGCCCGGCAAGAAGCUG CCGCUGGAGGUGCUCAAAGAGUUGGAAGCCAAUGCCCGGAAAGCUGGCUGCA CCAGGGGCUGUCUGAUCUGCCUGUCCCACAUCAAGUGCACGCCCAAGAUGAA GAAGUUCAUCCCAGGACGCUGCCACACCUACGAAGGCGACAAAGAGUCCGCA CAGGGCGGCAUAGGCGAGGCGAUCGUCGACAUUCCCCCU |
| dsRNA 400 bp | AGGGAGUCAAAGUUCUGUUUGCCCUGAUCUGCAUCGCUGUGGCCGAGGCCAA GCCCACCGAGAACAACGAAGACUUCAACAUCGUGGCCGUGGCCAGCAACUUC GCGACCACGGAUCUCGAUGCUGACCGCGGGAAGUUGCCCGGCAAGAAGCUG CCGCUGGAGGUGCUCAAAGAGUUGGAAGCCAAUGCCCGGAAAGCUGGCUGCA CCAGGGGCUGUCUGAUCUGCCUGUCCCACAUCAAGUGCACGCCCAAGAUGAA GAAGUUCAUCCCAGGACGCUGCCACACCUACGAAGGCGACAAAGAGUCCGCA CAGGGCGGCAUAGGCGAGGCGAUCGUCGACAUUCCUGAGAUUCCUGGGUUC AAGGACUUGGAGCCCUUGGAGCAGUUCAUCGCACCCCU |
| dsRNA 450 bp | AGGGAGUCAAAGUUCUGUUUGCCCUGAUCUGCAUCGCUGUGGCCGAGGCCAA GCCCACCGAGAACAACGAAGACUUCAACAUCGUGGCCGUGGCCAGCAACUUC GCGACCACGGAUCUCGAUGCUGACCGCGGGAAGUUGCCCGGCAAGAAGCUG CCGCUGGAGGUGCUCAAAGAGUUGGAAGCCAAUGCCCGGAAAGCUGGCUGCA CCAGGGGCUGUCUGAUCUGCCUGUCCCACAUCAAGUGCACGCCCAAGAUGAA GAAGUUCAUCCCAGGACGCUGCCACACCUACGAAGGCGACAAAGAGUCCGCA CAGGGCGGCAUAGGCGAGGCGAUCGUCGACAUUCCUGAGAUUCCUGGGUUC AAGGACUUGGAGCCCUUGGAGCAGUUCAUCGCACAGGUCGAUCUGUGUGUG GACUGCACAACUGGCUGCCUCAAAGGGCUUGCCCCCU |
| dsRNA 500 bp | AGGGAGUCAAAGUUCUGUUUGCCCUGAUCUGCAUCGCUGUGGCCGAGGCCAA GCCCACCGAGAACAACGAAGACUUCAACAUCGUGGCCGUGGCCAGCAACUUC GCGACCACGGAUCUCGAUGCUGACCGCGGGAAGUUGCCCGGCAAGAAGCUG CCGCUGGAGGUGCUCAAAGAGUUGGAAGCCAAUGCCCGGAAAGCUGGCUGCA |

| | |
|---|---|
| | CCAGGGGCUGUCUGAUCUGCCUGUCCCACAUCAAGUGCACGCCCAAGAUGAA<br>GAAGUUCAUCCCAGGACGCUGCCACACCUACGAAGGCGACAAAGAGUCCGCA<br>CAGGGCGGCAUAGGCGAGGCGAUCGUCGACAUUCCUGAGAUUCCUGGGUUC<br>AAGGACUUGGAGCCCUUGGAGCAGUUCAUCGCACAGGUCGAUCUGUGUGUG<br>GACUGCACAACUGGCUGCCUCAAAGGGCUUGCCAACGUGCAGUGUUCUGACC<br>UGCUCAAGAAGUGGCUGCCGCAACGCUGUGCCCCU |
| dsRNA<br>550 bp | AGGGAGUCAAAGUUCUGUUUGCCCUGAUCUGCAUCGCUGUGGCCGAGGCCAA<br>GCCCACCGAGAACAACGAAGACUUCAACAUCGUGGCCGUGGCCAGCAACUUC<br>GCGACCACGGAUCUCGAUGCUGACCGCGGGAAGUUGCCCGGCAAGAAGCUG<br>CCGCUGGAGGUGCUCAAAGAGUUGGAAGCCAAUGCCCGGAAAGCUGGCUGCA<br>CCAGGGGCUGUCUGAUCUGCCUGUCCCACAUCAAGUGCACGCCCAAGAUGAA<br>GAAGUUCAUCCCAGGACGCUGCCACACCUACGAAGGCGACAAAGAGUCCGCA<br>CAGGGCGGCAUAGGCGAGGCGAUCGUCGACAUUCCUGAGAUUCCUGGGUUC<br>AAGGACUUGGAGCCCUUGGAGCAGUUCAUCGCACAGGUCGAUCUGUGUGUG<br>GACUGCACAACUGGCUGCCUCAAAGGGCUUGCCAACGUGCAGUGUUCUGACC<br>UGCUCAAGAAGUGGCUGCCGCAACGCUGUGCGACCUUUGCCAGCAAGAUCCA<br>GGGCCAGGUGGACAAGAUCAAGGGGGCCGGUGGUGACCCU |
| dsRNA<br>1600 bp | AGGGCCGAGCGCAGAAGUGGUCCUGCAACUUUAUCCGCCUCCAUCCAGUCUA<br>UUAAUUGUUGCCGGGAAGCUAGAGUAAGUAGUUCGCCAGUUAAUAGUUUGCG<br>CAACGUUGUUGCCAUUGCUACAGGCAUCGUGGUGUCACGCUCGUCGUUUGG<br>UAUGGCUUCAUUCAGCUCCGGUUCCCAACGAUCAAGGCGAGUUACAUGAUCC<br>CCCAUGUUGUGCAAAAAAGCGGUUAGCUCCUUCGGUCCUCCGAUCGUUGUCA<br>GAAGUAAGUUGGCCGCAGUGUUAUCACUCAUGGUUAUGGCAGCACUGCAUAA<br>UUCUCUUACUGUCAUGCCAUCCGUAAGAUGCUUUUCUGUGACUGGUGAGUAC<br>UCAACCAAGUCAUUCUGAGAAUAGUGUAUGCGGCGACCGAGUUGCUCUUGCC<br>CGGCGUCAAUACGGGAUAAUACCGCGCCACAUAGCAGAACUUUAAAAGUGCU<br>CAUCAUUGGAAAACGUUCUUCGGGGCGAAAACUCUCAAGGAUCUUACCGCUG<br>UUGAGAUCCAGUUCGAUGUAACCCACUCGUGCACCCAACUGAUCUUCAGCAU<br>CUUUUACUUUCACCAGCGUUUCUGGGUGAGCAAAAACAGGAAGGCAAAAUGC<br>CGCAAAAAAGGGAAUAAGGGCGACACGGAAAUGUUGAAUACUCAUACUCUUCC<br>UUUUUCAAUAUUAUUGAAGCAUUUAUCAGGGUUAUUGUCUCAUGAGCGGAUA<br>CAUAUUUGAAUGUAUUUAGAAAAAUAAACAAAUAGGGGUUCCGCGCACAUUUC<br>CCCGAAAAGUGCCACCUGACGUCGACGGAUCGGGAGAUCUCCCGAUCCCCUA<br>UGGUGCACUCUCAGUACAAUCUGCUCUGAUGCCGCAUAGUUAAGCCAGUAUC<br>UGCUCCCUGCUUGUGUGUUGGAGGUCGCUGAGUAGUGCGCGAGCAAAAUUU<br>AAGCUACAACAAGGCAAGGCUUGACCGACAAUUGCAUGAAGAAUCUGCUUAG<br>GGUUAGGCGUUUUGCGCUGCUUCGCGAUGUACGGGCCAGAUAUACGCGUUG<br>ACAUUGAUUAUUGACUAGUUAUUAAUAGUAAUCAAUUACGGGGUCAUUAGUUC<br>AUAGCCCAUAUAUGGAGUUCCGCGUUACAUAACUUACGGUAAAUGGCCCGCC<br>UGGCUGACCGCCCAACGACCCCCGCCCAUUGACGUCAAUAAUGACGUAUGUU<br>CCCAUAGUAACGCCAAUAGGGACUUUCCAUUGACGUCAAUGGGUGGAGUAUU<br>UACGGUAAACUGCCCACUUGGCAGUACAUCAAGUGUAUCAUAUGCCAAGUAC<br>GCCCCCUAUUGACGUCAAUGACGGUAAAUGGCCCGCCUGGCAUUAUGCCCAG<br>UACAUGACCUUAUGGGACUUUCCUACUUGGCAGUACAUCUACGUAUUAGUCA<br>UCGCUAUUACCAUGGUGAUGCGGUUUUGGCAGUACAUCAAUGGGCGUGGAUA<br>GCGGUUUGACUCACGGGGAUUUCCAAGUCUCCACCCCAUUGACGUCAAUGGG<br>AGUUUGUUUUGGCACCAAAAUCAACGGGACUUUCCAAAAUGUCGUAACAACUC<br>CGCCCCAUUGACGCAAAUGGGCGGUAGGCGUGUACGGUGGGAGGUCUAUAU<br>AAGCAGAGCUCUCCCU |

**Dataset S1 (separate file).** Data used to prepare Fig. S7-S10.
